# Projected habitat loss and spatial redistribution of two alpine and subalpine species in Himachal Pradesh, Western Himalaya under CMIP6 scenarios

**DOI:** 10.64898/2026.09.20.752453

**Authors:** Simran Tomar, Merja Helena Tölle, Matthijs Vos

**Affiliations:** Institute of Water, Waste and Environmental Engineering, University of Kassel, 34125 Kassel, Germany; Theoretical and Applied Biodiversity Research, Faculty of Biology and Biotechnology, Ruhr University Bochum, Bochum, Germany

**Keywords:** Species distribution modelling, Habitat suitability, Climate change, *Meconopsis aculeata*, *Rhododendron campanulatum*, Conservation planning

## Abstract

Climate change will alter habitat suitability for plant communities in mountain ecosystems of the Western Himalaya. There will be contrasting magnitudes and spatial configurations of these changes for different plant species, and it is an important question how to translate these contrasts into effective climate-proof multi-species conservation planning. This study assessed current habitat associations and projected changes in suitable habitat for two contrasting species, *Meconopsis aculeata* and *Rhododendron campanulatum*, in Himachal Pradesh, India. Field surveys and spatially filtered occurrence records were combined with MaxEnt modelling using 10 non-collinear environmental predictors. Habitat suitability was projected to 2050 and 2070 under Shared Socioeconomic Pathways; SSP1-2.6, SSP2-4.5 and SSP5-8.5 using HadGEM3-GC31-LL.

Field observations showed contrasting habitat associations, with *M. aculeata* occurring mainly in open alpine herbaceous communities and *R. campanulatum* in subalpine woody communities. Elevation was an important predictor for both species while climatic predictors contributed differently to their relative importance and response patterns. Future projections indicated reductions in suitable habitat of approximately 25% for *M. aculeata* and 27-28% for *R. campanulatum*. Habitat-change maps showed heterogeneous patterns of loss, gain and persistence rather than uniform contraction, and centroid displacement remained small (7.7-8.5 km). We performed proximity analyses to show which currently suitable areas are most close to areas of future gain, in order to support the design of climate-proof protected areas. We discuss how interspecific differences in response are relevant to multi-species conservation planning that considers the spatial relationship between current high-suitability areas and the gain, loss and redistribution of future suitable habitat.

## 1. Introduction

Alpine ecosystems are among the most climate-sensitive terrestrial environments because species occupying high elevations often exhibit narrow thermal tolerances, short growing seasons, specialised habitat requirements, and limited opportunities for dispersal. Consequently, even modest increases in temperature can alter species distributions, disrupt community composition, and reduce ecosystem resilience (Körner 2003; Pepin et al. 2022; Sekar et al. 2025). Mountain biodiversity hotspots are therefore expected to experience some of the most pronounced biological responses to climate change (Ohlemüller et al. 2008), making them natural laboratories for the study of climate-driven species redistribution.

The Himalayan region, recognised as a global biodiversity hotspot (Myers et al. 2000), supports a wealth of unique and endemic flora increasingly threatened by climate change (Kattel 2022; Ahmad et al. 2025) . *Meconopsis aculeata* Royle, a rare Himalayan blue poppy and *Rhododendron campanulatum* D. Don, a dominant alpine shrub, hold significant ecological and cultural importance in Himachal Pradesh (Basnett and Ganesan 2022; Manzoor et al. 2025). They occupy contrasting altitudinal ranges and have alternative habitat preferences (Shaheen et al. 2023). The two species also represent contrasting life forms- a perennial alpine herb (*M. aculeata*) and a dominant evergreen subalpine shrub (*R. campanulatum*). The above suggests they could serve as complementary indicator species for the consequences of climate change.

Species responses to environmental change can provide useful insights into how individual components of alpine ecosystems may respond to changing environmental conditions. In particular, species with restricted distributions or strong associations with particular environmental conditions can reveal spatial patterns of habitat change and help identify areas where suitable conditions may persist or emerge under future scenarios. However, responses are likely to differ among species because the environmental variables governing their distributions may differ.

*M. aculeata* Royle (family Papaveraceae) is a deciduous perennial herb primarily distributed at elevations of 3,300 - 4,600 m. It is a rare medicinal plant with a largely western Himalayan distribution, although isolated occurrences have also been reported from the eastern Himalayas (Paul et al., 2024). The species typically inhabits moist, shaded alpine grasslands, stream banks, and rock crevices with semi-arid to mesic soils (Hooker 1890; Majid et al. 2015). The species is characterised by conspicuous blue flowers borne on prickly stems, producing numerous seeds within dehiscent capsules (Ganaie et al., 2016; Shukla et al., 2021) **(Fig. S1a).**

In contrast, *R. campanulatum* (family Ericaceae) is an evergreen, gregarious shrub distributed more widely across the sub-alpine and outer alpine ranges of the Himalayas, from Kashmir to Bhutan, at elevations between 9,000 and 14,000 ft. (Kala 2003). Though more prevalent than *M. aculeata*, it remains sensitive to extreme climatic fluctuations such as frost and drought (Singh et al. 2025). *R. campanulatum* often forms dense krummholz stands that strongly influence subalpine vegetation structure, snow accumulation, soil temperature and regeneration dynamics, making the species an important structural component of Himalayan treeline ecosystems (Kumar and Khanduri 2024) **(Fig. S1b)**.

*M. aculeata* is associated with specialised alpine meadow communities, whereas *R. campanulatum* dominates upper subalpine shrublands and krummholz vegetation. These contrasting community associations reflect distinct ecological niches. This suggests a potential suitability as complementary indicators of environmental change across the alpine-subalpine transition (Majid et al. 2015; Shukla et al. 2021; Kumar and Khanduri 2024; Manzoor et al. 2025). Both species possess considerable ethnobotanical importance and are widely used in traditional Himalayan medicine and local livelihoods **(Online Resource 1, Table S1).**

Species distribution models (SDMs) have become widely used for assessing potential changes in species distributions under environmental change and for identifying areas where suitable environmental conditions may persist or emerge under future scenarios (Elith and Leathwick 2009; Franklin 2010). MaxEnt is one of the most widely used presence-only approaches for species distribution modelling and is particularly suitable when occurrence data are limited (Phillips et al. 2006; Elith and Leathwick 2009; Booth 2022). Future projections can be evaluated under different Shared Socioeconomic Pathways (SSPs), which represent alternative trajectories of greenhouse-gas emissions and associated climate change. SSP1-2.6 represents a low-emissions pathway, SSP2-4.5 an intermediate pathway, and SSP5-8.5 a high-emissions pathway, thereby providing a range of plausible future climate conditions. For species distribution modelling, the relevance of these scenarios lies in their projected changes in climatic conditions, particularly temperature- and precipitation-related variables, which can alter the environmental conditions associated with species’ current habitat suitability. Comparing these pathways therefore allows assessment of whether and how projected changes in climatic conditions may alter the spatial distribution of suitable habitat over time.

Previous studies have projected potential range shifts for Himalayan alpine plants, including *Meconopsis* and *Rhododendron*, under future climate scenarios (Basnett and Ganesan, 2022; Dad and Rashid, 2022; Kumar and Khanduri, 2024; Paul and Samant, 2024). However, most studies have focused on individual species in particular areas and relied primarily on climatic predictors. Field-based ecological and phytosociological assessments are rarely integrated with spatial modelling to compare species-specific responses and evaluate their implications for regional climate-proof conservation planning. Consequently, important knowledge gaps remain regarding how habitat suitability, connectivity and spatial distribution change across contrasting species and climate scenarios, how much of the currently suitable habitat remains suitable in the future, and whether areas identified for one species also retain conservation relevance for other species with partly different ecological requirements. These questions are particularly relevant because the responses of individual species to climate change may vary among landscapes and because no single species can be expected to represent the habitat requirements of the wider alpine flora. This study integrates field-based ecological surveys, phytosociological observations, species distribution modelling, and future CMIP6 climate projections to assess two ecologically contrasting alpine species and compare their projected habitat responses. By considering the two species together, the study further explores their potential complementary value for conservation planning rather than assuming that either species alone can serve as a universal indicator of climate-driven change. The analysis identifies areas of present and future habitat suitability and shows where these are in close proximity, for each of the two species. We aim to show how this supports climate-proof conservation planning and the design of protected areas for species that may face dispersal limitation under climate-induced redistribution. Thus, the present study aims to:

a. conduct an ecological assessment of *M. aculeata* and *R. campanulatum* in their natural habitats to characterize their current distribution and associated plant communities;
b. evaluate how suitable habitats for *M. aculeata* and *R. campanulatum* may change under present and future climate scenarios using the MaxEnt modelling approach;
c. quantify projected habitat redistribution, including gain, loss, and stable areas, together with centroid displacement and changes in similarity between current and future suitable-habitat distributions under alternative CMIP6 scenarios; and
d. assess the potential complementary value of *M. aculeata* and *R. campanulatum* as ecological indicators by comparing their species-specific habitat responses and identifying the extent to which conservation priorities based on either species alone may differ from those emerging from their combined consideration.

The findings provide species-specific evidence to support climate-sensitive biodiversity conservation and nature reserve design. Integrating these findings with studies on a wider range of species will help policymakers and forest managers identify areas that are suitable in the present that are well-connected to areas that are most likely highly suitable in the future, for ensembles of multiple species. We discuss how our results are on the one hand specific to the Himachal Pradesh area and on the other hand have much wider implications for conservation, nature reserve planning and the use of indicator species.

## 2. Material and methods

### 2.1 Field surveys and population assessment

Population assessments of *M. aculeata* and *R. campanulatum* were conducted in the subalpine and alpine regions of Himachal Pradesh between 2018 and 2024 during the peak flowering season. Field surveys recorded *M. aculeata* at two sites and *R. campanulatum* at twelve sites. These field observations were used to assess population characteristics and ecological attributes and also contributed verified occurrence records for species distribution modelling. Population characteristics were estimated using a stratified random sampling approach along the elevational gradient. Quadrat sampling was carried out using 1 × 1 m quadrats for the herbaceous *M. aculeata* and 5 × 5 m quadrats for the shrub *R. campanulatum*. Twenty quadrats were established at each site, and the number of individuals was recorded following standard ecological methods. Frequency (%) was calculated as the number of quadrats in which a species occurred divided by the total number of quadrats sampled, multiplied by 100. Abundance was calculated as the total number of individuals of a species divided by the number of quadrats in which that species occurred and was expressed as individuals per occupied quadrat. Herb density was expressed as individuals m⁻², whereas shrub density was standardized to individuals ha⁻¹ to facilitate comparison among shrub populations following standard vegetation sampling procedures (Misra 1968) **(Fig. 1).** Sampling locations and associated metadata are provided in **Online Resource 1, Table S2.**

**Fig. 1.**
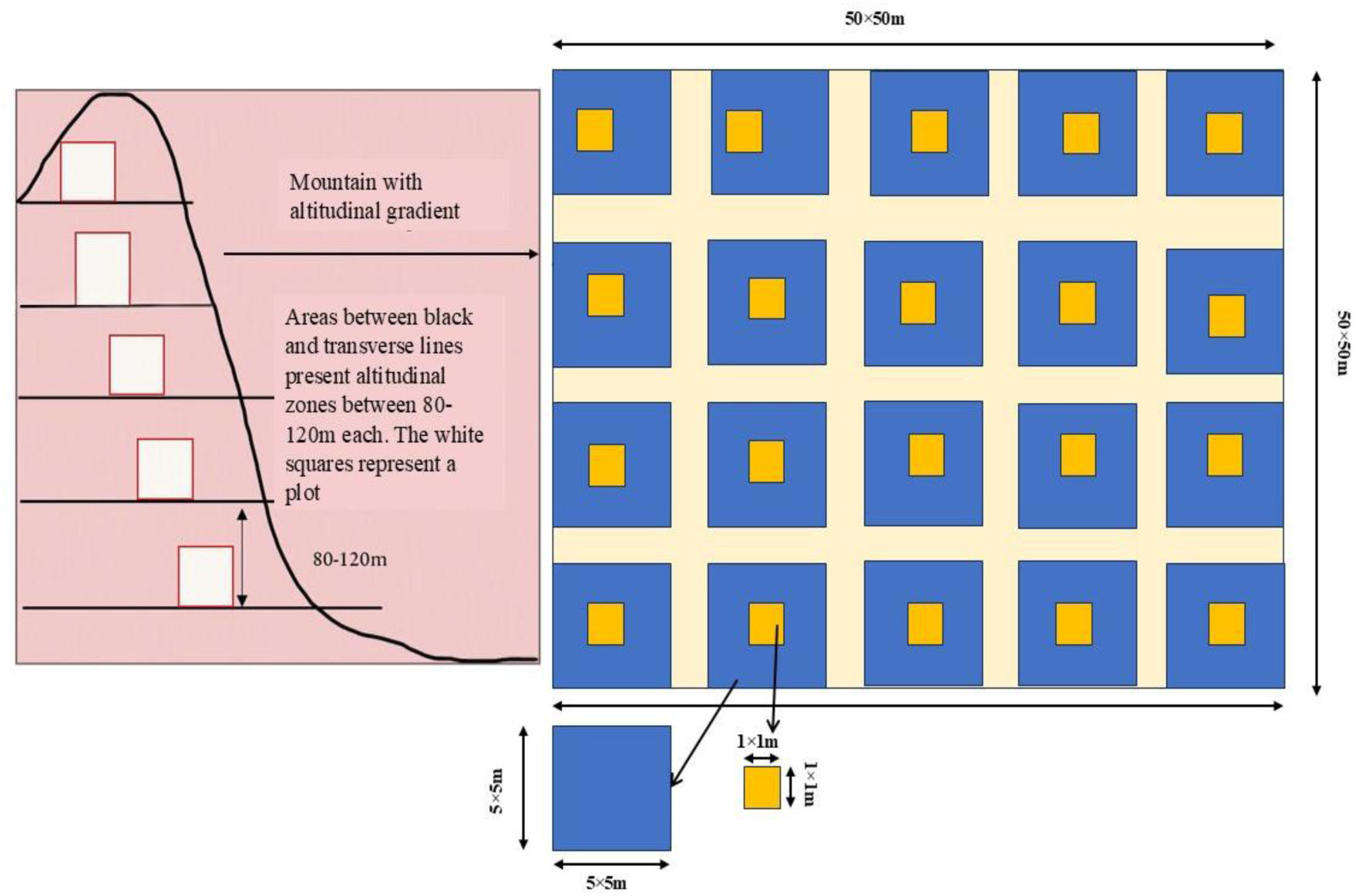
Design of sampling plot for population assessment

### 2.2 Species data

Occurrence records for *M. aculeata* and *R. campanulatum* were compiled from multiple sources, including field surveys conducted during this study, published literature, theses, online herbarium databases, national herbarium records, and the Global Biodiversity Information Facility (GBIF; https://www.gbif.org), **(Fig. 2)**. Duplicate records, records lacking geographic coordinates, obvious spatial errors, and records with inconsistent locality information were removed during data cleaning. Coordinates were subsequently verified using Google Earth Pro (v7.3.6.9796) and ArcGIS to ensure spatial accuracy. Using R (version 4.2.2; R Core Team, 2022), spatial thinning was performed using a 5 × 5 km grid with the *spThin* package implemented through the Wallace platform (v2.0.5) to reduce spatial sampling bias and autocorrelation among occurrence records (Chauhan et al. 2022; Johnson et al.). After filtering, 68 occurrence records for *M. aculeata* and 65 records for *R. campanulatum* were retained for species distribution modelling. The occurrence dataset, including geographic coordinates, is provided in **Online Resource 1, Table S3.**

**Fig. 2.**
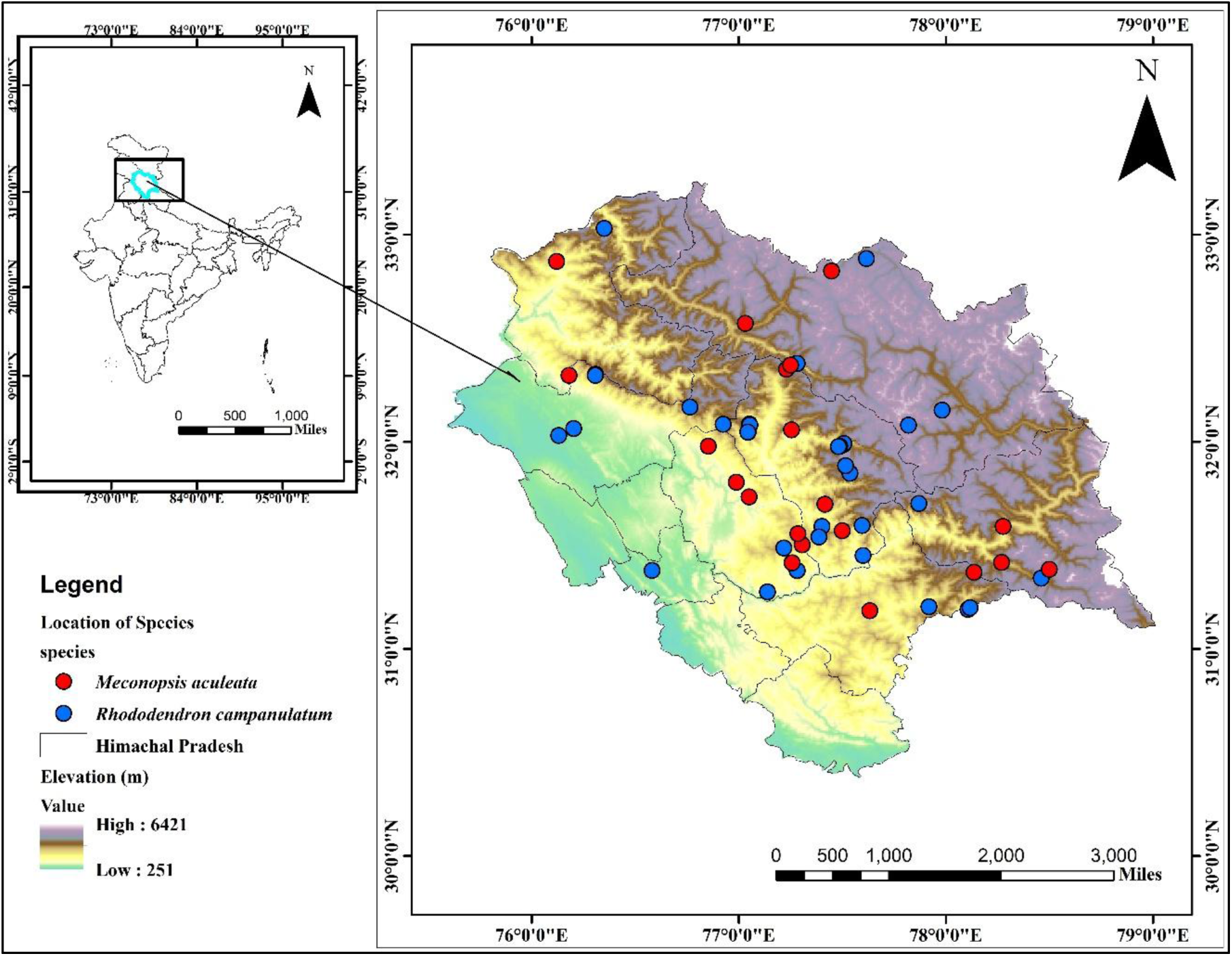
Map of study area showing the primary locations of *M. aculeata* (red dots) and *R. campanulatum* (blue dots) in Himachal Pradesh along with the elevation profile.

### 2.3 Climatic data and other environmental variables

Environmental predictors were selected to represent the major climatic, topographic, and edaphic factors known to influence the distribution of alpine and subalpine plant species. Climatic variables were obtained from the WorldClim version 2.1 database (Fick and Hijmans 2017) including 19 bioclimatic variables derived from monthly temperature and precipitation data at a spatial resolution of 30 arc-seconds (∼1 km). Elevation data were obtained from the Shuttle Radar Topography Mission (SRTM), while slope and aspect were derived from the digital elevation model using ArcGIS. Land-cover data were obtained using the R package *geodata* and represented categorical land-cover classes, whereas soil data were obtained from the Harmonized World Soil Database (HWSD v1.2), with soil classes based on the World

Reference Base for Soil Resources (WRB 2014, update 2015), available through the Food and Agriculture Organization (FAO) soil portal. Land-cover and soil datasets were resampled to match the 30 arc-second resolution of the climatic variables prior to analysis. The numerical values assigned to land-cover and soil classes represent categorical codes and therefore should not be interpreted as continuous environmental gradients. To minimise multicollinearity among predictor variables, Pearson’s correlation coefficients were calculated and highly correlated variables (|r| > 0.70) were excluded from further analyses. Variance Inflation Factor (VIF) analysis was subsequently used to confirm the independence of the retained predictors. Ten variables with low multicollinearity and high ecological relevance were retained for model development **(Online Resource 1, Table S4 and S5).** Because topographic variables (elevation, slope, and aspect) remain relatively stable over the temporal scale of climate projections, the same topographic layers were used for both current and future models.

### 2.4 Model design and Species Distribution modelling

Future habitat suitability was projected for 2050 (2041-2060) and 2070 (2061-2080) using the HadGEM3-GC31-LL Global Climate Model from the Coupled Model Intercomparison Project Phase 6 (CMIP6). Three Shared Socioeconomic Pathways (SSP1-2.6, SSP2-4.5, and SSP5- 8.5), representing low-, intermediate-, and high-emission scenarios, respectively, were used to evaluate the potential impacts of climate change on species distributions (O’Neill et al., 2017; IPCC, 2023). Future climate layers were obtained from statistically downscaled and bias-corrected WorldClim version 2.1 CMIP6 projections at 30 arc-second (∼1 km) spatial resolution (Fick and Hijmans 2017). The HadGEM3-GC31-LL model was selected because it provides high-resolution CMIP6 climate projections and has been widely applied in studies of Himalayan climate and species distribution modelling (Kumar and Sarthi 2021; Meher and Das 2024; Tiwary et al. 2024).

Species distributions were modelled using MaxEnt version 3.4.4k (Phillips et al. 2006), a machine-learning algorithm designed for presence-only data that performs well when modelling rare species with limited occurrence records (Phillips et al. 2006; Merow et al. 2013; Phillips et al. 2022). Models were generated using ten replicate runs with cross-validation. Occurrence records were randomly partitioned into training (75%) and testing (25%) datasets, and each replicate was evaluated using an independent test dataset. The default feature classes in MaxEnt were retained because they provide robust performance for moderate sample sizes while reducing unnecessary model complexity. Model settings included 10,000 background points, a regularization multiplier of 1.0, and a maximum of 500 iterations to reduce overfitting while maintaining model generality.

Model performance was evaluated using the Area Under the Receiver Operating Characteristic Curve (AUC) calculated from the independent 25% test dataset. AUC values range from 0.5 (random prediction) to 1.0 (perfect discrimination), with higher values indicating better predictive performance (Swets 1988; Allouche et al. 2006). Response curves, receiver operating characteristic (ROC) curves, and jackknife analyses used to assess variable importance are presented in the **Online Resource 2, Figures S2-S4.**

### 2.5 Future range shift and centroid analysis

To quantify potential spatial shifts in habitat suitability under climate change, centroid analyses were conducted using habitat suitability projections generated by MaxEnt. For each species, the geographic centroid of the predicted suitable habitat was calculated for the current distribution and future projections under SSP1-2.6, SSP2-4.5, and SSP5-8.5 for 2050 (2041- 2060) and 2070 (2061-2080). The centroid therefore represents the geographic centre of the predicted suitable habitat rather than the geometric centre of the species’ distribution. Range-shift metrics included: (i) centroid displacement (km), calculated as the straight-line distance between current and future centroids; (ii) bearing (degrees), representing the direction of centroid movement relative to geographic north; and (iii) Schoener’s D, which quantifies niche overlap between current and future habitat suitability projections, ranging from 0 (no overlap) to 1 (complete overlap) (Warren et al. 2008). All analyses were performed in **R (version 4.2.2; R Core Team, 2022).**

## 3. Results

### 3.1 Distribution, Population structure and habitat preference

Field surveys recorded *M. aculeata* at two natural populations between 3,264 and 4,176 m above sea level whereas *R. campanulatum* was documented at twelve field populations between 2,850 and 3,851 m a.s.l. in Himachal Pradesh **(Online Resource 1, Table S2).** *M. aculeata* occurred as small, localized populations with low frequency (10-20%) and mean abundance (2.00-2.25 individuals per occupied quadrat), whereas *R. campanulatum* formed widespread subalpine stands with substantially higher frequency (35-100%) and mean abundance (1.46- 30.15 individuals per occupied quadrat) **(Table 1).** Habitat surveys recorded 67 associated plant species belonging to 56 genera and 25 families within *M. aculeata* habitats, whereas *R. campanulatum* habitats supported 156 associated species representing 107 genera and 47 families, indicating considerably greater floristic diversity within subalpine shrub communities.

**Table 1.** Phytosociological characteristics of *M. aculeata* and *R. campanulatum* populations in Himachal Pradesh.

| Species | Density <sup>1</sup> | Diversity (H') | Frequency (%) | Abundance | A/F ratio |
| --- | --- | --- | --- | --- | --- |
| *M. aculeata* | 0.2 | 0.014 | 10.00 | 2.0 | 0.2 |
|  | 0.45 | 0.022 | 20.00 | 2.25 | 0.11 |
| *R. campanulatum* | 110 | 0.21 | 75.00 | 1.46 | 0.01 |
|  | 65 | 0.18 | 35 | 1.85 | 0.05 |
|  | 230 | 0.34 | 65 | 3.53 | 0.05 |

<sup>1</sup> Density is expressed as individuals m<sup>-2</sup> for herbaceous communities (*M. aculeata*) and Individuals ha<sup>-1</sup> for shrub communities (*R. campanulatum*), reflecting standard phytosociological sampling methods for different vegetation strata. Abundance is expressed as Individuals per occupied quadrat for both species.

|  |  |  |  |  |  |
| --- | --- | --- | --- | --- | --- |
|  | 365 | 0.35 | 100 | 3.65 | 0.03 |
|  | 115 | 0.24 | 60 | 1.91 | 0.03 |
|  | 110 | 0.36 | 85 | 3.23 | 0.03 |
|  | 150 | 0.32 | 55 | 2.72 | 0.04 |
|  | 185 | 0.35 | 65 | 2.84 | 0.04 |
|  | 195 | 0.44 | 55 | 3.54 | 0.06 |
|  | 3015 | 0.35 | 100 | 30.15 | 0.30 |
|  | 1235 | 0.31 | 100 | 12.35 | 0.12 |
|  | 1235 | 0.34 | 100 | 12.35 | 0.12 |

Vegetation structure differed markedly between the two habitats*. Rhododendron campanulatum* communities were dominated by dense shrub vegetation, including *Rosa macrophylla*, *Cotoneaster microphyllus*, *Berberis aristata*, *Juniperus indica*, and *Rhododendron lepidotum*. In contrast, *M. aculeata* occurred in open alpine herbaceous communities characterised by species such as *Polygonum paronychioides*, *Rhodiola quadrifida*, *Potentilla fulgens*, and *Anaphalis triplinervis*. These contrasting assemblages reflect distinct ecological niches across the alpine-subalpine transition.

The observed contrast in vegetation structure reflects the altitudinal transition from shrub-dominated subalpine ecosystems to herb-dominated alpine communities, illustrating the contrasting ecological requirements of the two focal species. Only 14 species belonged both to the group of 67 *M. aculeata*-associated species and to the 156 *R. campanulatum*-associated species. Importantly, due to contrasting preference for habitat types, *M. aculeata* and *R. campanulatum* were never associated with each other in the surveyed sites.

Across all associated taxa, herbs contributed the greatest species richness in both habitats, accounting for 94% of the associated flora in *M. aculeata* habitats and 78% in *R. campanulatum* habitats. Shrubs comprised only 6% of the flora associated with *M. aculeata* but increased to 14% in *R. campanulatum* communities, while trees represented 6% of associated species in the latter. Among the recorded families, Asteraceae exhibited the greatest species richness, followed by Rosaceae, Polygonaceae, and Ranunculaceae. The complete list of associated plant species and community types for each site is provided in **Online Resource 1, Table S6.**

### 3.2 Environmental predictors of current habitat suitability

Jackknife analyses of regularized training gain indicated that the relative importance of environmental predictors differed between the two species. For *M. aculeata*, elevation provided the highest gain when used individually (approximately 0.30), followed by BIO15 (Precipitation Seasonality, 0.21), BIO8 (Mean Temperature of Wettest Quarter, 0.17), BIO19 (Precipitation of Coldest Quarter, 0.14), and BIO2 (Mean Diurnal Temperature Range, 0.11). Soil (0.06), slope (0.07), BIO3 (Isothermality, 0.04), land cover (0.03), and aspect (0.01) provided comparatively lower gains when considered individually.

For *R. campanulatum*, elevation also provided the highest gain when used individually (approximately 0.98), followed by BIO2 (0.92), land cover (0.56), BIO15 (0.51), BIO19 (0.48), BIO8 (0.42), BIO3 (0.38), slope (0.17), aspect (0.13), and soil (0.02). The exclusion of individual variables produced different reductions in regularized training gain, indicating that the predictors contributed differently to model performance. Overall, elevation was the strongest individual predictor for both species, while the relative contributions of the climatic and other environmental variables differed between the models **(Fig. 3).**

**Fig. 3.**
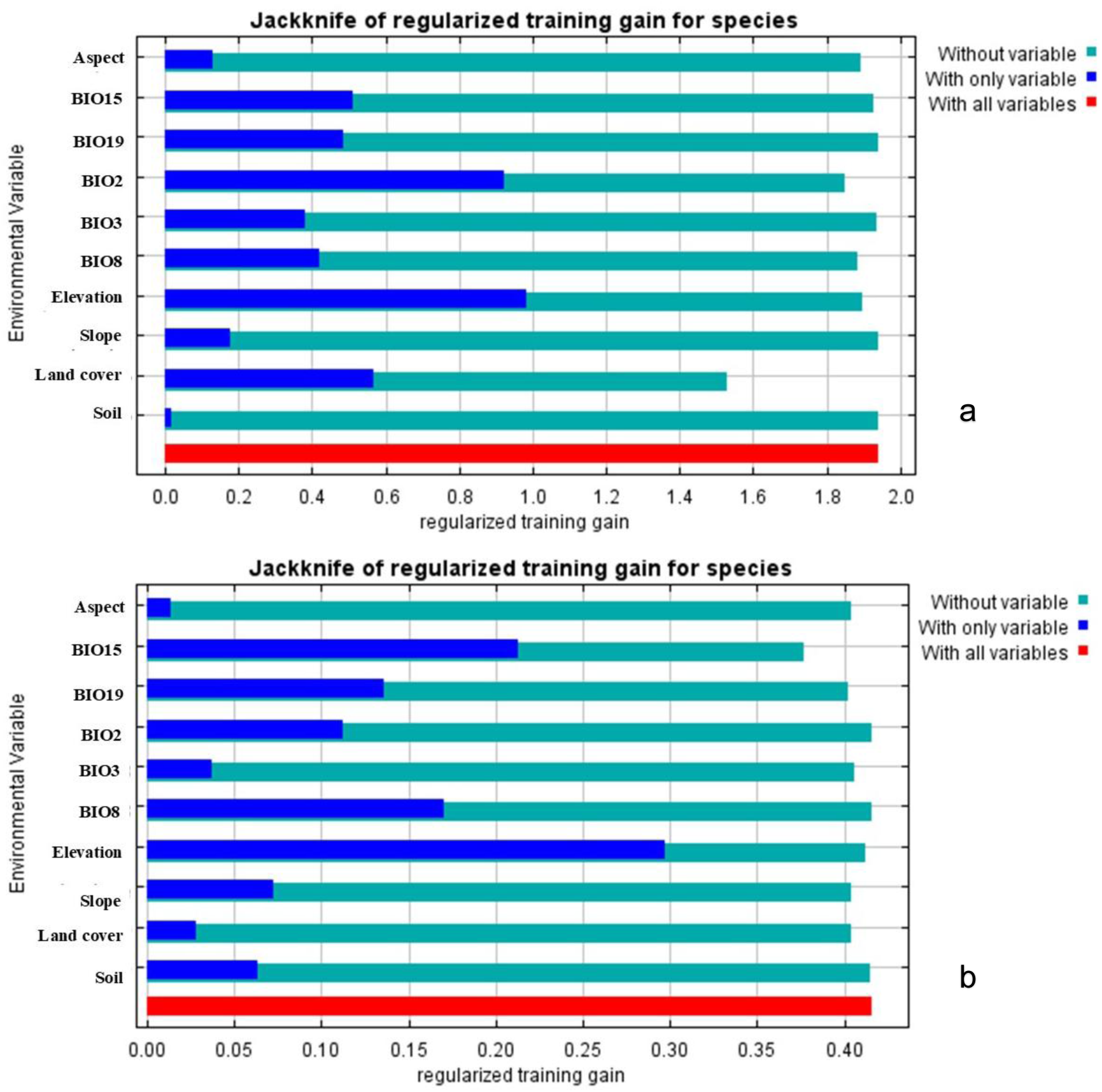
Result of jackknife test for evaluating the relative contribution of the predictor environmental variables to the habitat model of (a) *R. campanulatum* and (b) *M. aculeata*; Abbreviations used BIO8 = Mean Temperature of Wettest Quarter; BIO3 = Isothermality; BIO2= mean diurnal range; BIO19 = Precipitation of Coldest Quarter; BIO15= Precipitation Seasonality

### 3.3 Species responses to key environmental gradients

Full-model MaxEnt response curves showed distinct but partially overlapping patterns in predicted suitability across the environmental predictors for *M. aculeata* and *R. campanulatum* **(Online Resource 2, Figs. S2 and S3).** The y-axis represents the MaxEnt logistic output (predicted habitat suitability), whereas the x-axis represents the value or coded class of the respective environmental predictor.

For *M. aculeata*, predicted suitability increased with elevation up to approximately 2,200 m and remained relatively high up to 2,500m above sea level, after which the response declined gradually at higher elevations. Suitability increased rapidly with slope and remained relatively stable at values above 20°. The species response to aspect was relatively stable up to approximately 140°, followed by a gradual decline toward higher aspect values. Among climatic predictors, predicted suitability increased with BIO3 (isothermality) from approximately 27 to 33, after which the response approached a plateau and a gradual decline at values higher than 35. In contrast, suitability declined progressively with increasing BIO15 (precipitation seasonality), particularly above approximately 40mm. The response to BIO19 (precipitation of the coldest quarter) increased gradually up to approximately 220 mm before declining at higher values. Suitability also declined progressively with increasing BIO2 (mean diurnal temperature range), while BIO8 (mean temperature of the wettest quarter) showed stable predicted suitability up to 5° C and then showed a pronounced peak at approximately 17° C and declined sharply at higher temperatures.

For *R. campanulatum*, predicted suitability increased strongly with elevation up to approximately 3,000 m, reached a pronounced maximum at approximately 3,200m, and subsequently declined at higher elevations. Suitability increased rapidly at lower slope values, up to 50° and subsequently remained relatively stable across higher slopes. The response to aspect declined gradually up to 180°, and subsequently stable from 200 - 320°, followed by a marked increase at high aspect values up to 360° and stable at higher values than 360°. Among climatic predictors, predicted suitability declined strongly with increasing BIO2, (mean diurnal temperature range), particularly between approximately 12 and 14°C. For BIO3 (isothermality), suitability increased gradually to approximately 33.5 and remained stable up to 34 and subsequently declined at higher values. The response to BIO8 (mean temperature of the wettest quarter) showed a more complex patter with two pronounced peaks at -7.5° C and 14° C and stable predicted suitability from 1°-5° C. For BIO15 (precipitation seasonality), suitability declined with increasing precipitation values. The response curve for BIO19 (precipitation of the coldest quarter) peaked at 220m before declining at higher precipitation values.

For land cover, both species showed step-like changes in predicted suitability across the coded land-cover classes. For both *M. aculeata* and *R. campanulatum*, predicted suitability increased after 0.0 at the lower coded values and subsequently remained relatively stable across higher values up to 0.79 and 0.55. However, the predicted habitat suitability declined at higher values up to 1.0 for both species, while showing a sharp peak at 1.0 for *M. aculeata*. Because land cover represents categorical classes, these numerical codes do not represent a continuous ecological gradient. Similarly, the response to soil type differed among coded soil classes rather than representing a continuous response to increasing soil values. For *M. aculeata*, predicted suitability was relatively high at lower coded soil values, followed by a peak decline at coded value 4,000 and relatively stable values thereafter. For *R. campanulatum*, predicted suitability increased across the lower-to-intermediate coded soil classes and subsequently approached a plateau at higher coded values.

Overall, the response curves indicate that elevation was an important predictor for both species, while both temperature and precipitation related variables show similar structured relationships with predicted suitability. The two species differed in the relative importance and shape of individual predictor responses, but these differences did not indicate a clear separation between precipitation-related and temperature-related controls. Instead, the results suggest that habitat suitability in both species was associated with different combinations of thermal variability, seasonal temperature conditions, precipitation seasonality, and precipitation during the cold season. The top-five of most important predictors was similar for both species, including elevation, BIO2, BIO15 and BIO19 in both cases (Fig. 3). The exact response curves for these predictors varied between the two species as described above (Figs. S2, S3). A further contrast was that BIO8 was included in the top-five of *M. aculeata*, whereas this was landcover for *R. campanulatum* (Fig. 3).

### 3.4 Current and projected habitat suitability

The MaxEnt models showed good predictive performance for both species, with AUC values of 0.830 for *M. aculeata* and 0.976 for *R. campanulatum*, respectively **(Online Resource 2, Fig. S4).**

Under current climatic conditions, suitable habitats for both species were concentrated mainly in the alpine and subalpine regions of Himachal Pradesh (Fig. 5). For *M. aculeata*, medium- and high-suitability habitats accounted for 26.71% and 31.63% of the total suitable area, respectively. For *R. campanulatum*, the corresponding proportions were 20.03% and 19.96% **(Online Resource 2, Table S7).**

The current distribution maps showed different spatial patterns between the two species. Highly suitable areas for *M. aculeata* were distributed across high-elevation regions of Kinnaur, Kullu, Chamba, Shimla, and Lahaul-Spiti, whereas suitable areas for *R. campanulatum* were more spatially restricted and concentrated mainly in high-elevation areas of Chamba, Kullu, Mandi, and Shimla **(Fig. 4).**

**Fig. 4.**
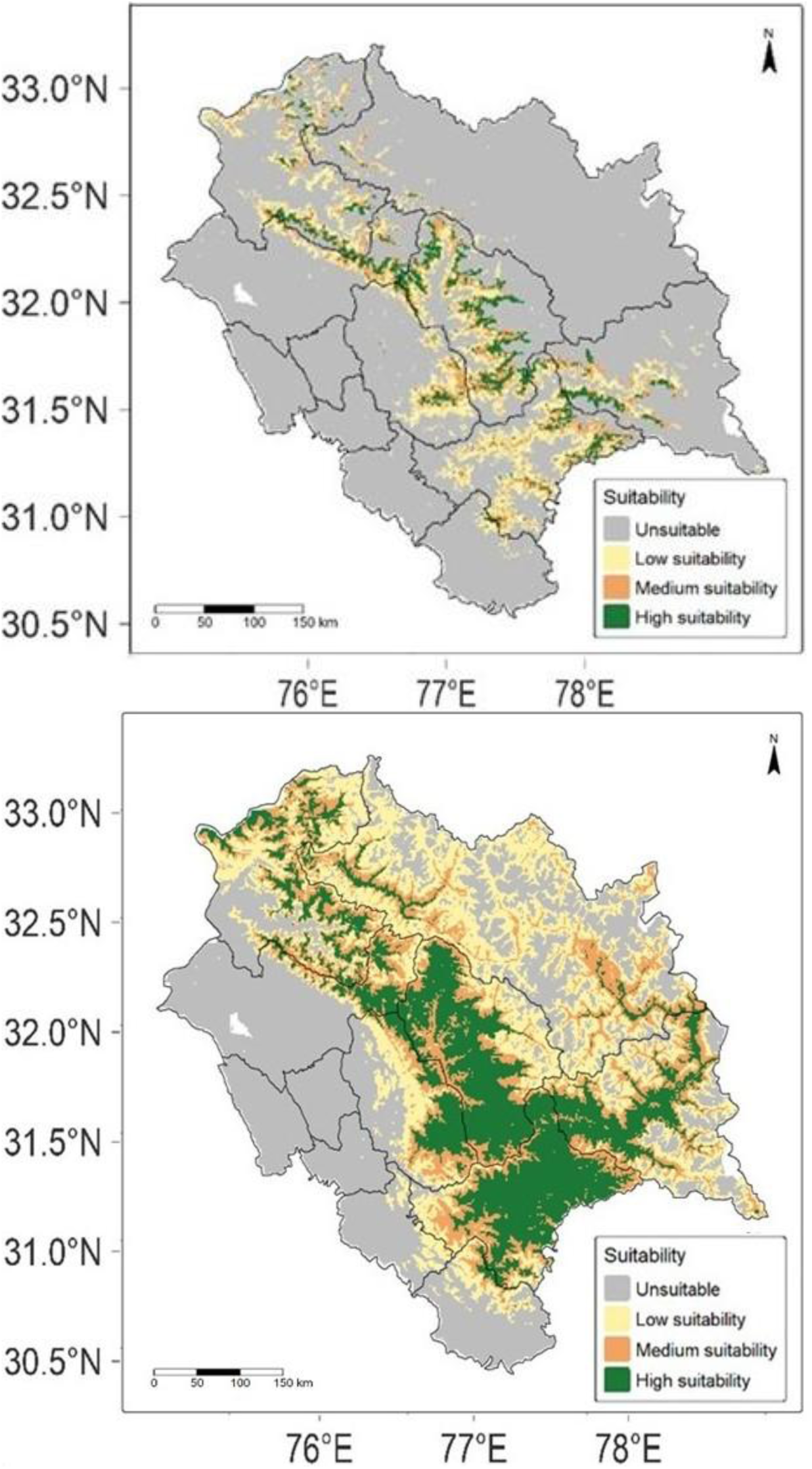
Current habitat suitability of *M. aculeata* (top panel) and *R. campanulatum* (bottom panel) Maxent 3.4.3 (https://biodiversityinformatics.amnh.org/open_source/maxent/); ArcMap 10.8 (https://unikassel-its.maps.arcgis.com/home/)

Future projections indicated a reduction in the total area classified as suitable for *M. aculeata*under all scenarios and periods (Table 2). The current suitable area of 2,785.82 km² decreased to 2,076.43 km² under SSP1-2.6 in 2050, corresponding to a 25.46% reduction. By 2070, suitable area ranged from 2,067.87 km² under SSP1-2.6 to 2,090.09 km² under SSP5-8.5, corresponding to reductions of 25.77% and 24.97%, respectively. The differences among scenarios were therefore relatively small, with reductions of approximately 25% across all projections **(Online Resource 2, Fig. S5).**

For *R. campanulatum*, the current suitable area of 777.89 km² decreased under all future scenarios (Online Resource Table S6). Projected suitable area ranged from 558.05 km² under SSP2-4.5 in 2070 to 564.75 km² under SSP1-2.6 in 2070. Under SSP5-8.5, suitable area decreased to 563.45 km² in 2050 and 557.86 km² in 2070, corresponding to reductions of 27.56% and 28.28%, respectively **(Online Resource 2, Fig. S6)**.

The suitability-change maps showed that both species experienced spatially heterogeneous changes between current and projected conditions, comprising areas of suitability loss, stability, and gain **(Online Resource 3, Fig. S7-S8)**. For *M. aculeata*, areas of both gain and loss were distributed across the species’ current and projected suitable range, with stable areas also persisting across substantial portions of the distribution. Similar spatially heterogeneous patterns were observed for *R. campanulatum*, although the areas of suitability change were more spatially concentrated within its comparatively restricted distribution. These maps therefore show that the projected reduction in total suitable area does not represent uniform loss across the entire landscape; rather, loss occurs while suitable and unsuitable areas are spatially redistributed under future climate conditions. For *M. aculeata*, future scenarios show a clear band of gain mostly south-west of currently suitable areas. For *R. campanulatum*, gain areas were relatively small and occurred mostly north of presently suitable habitat, mainly on the north-east and north-west ends of the present distribution **(Fig. 5)**.

**Fig. 5.**
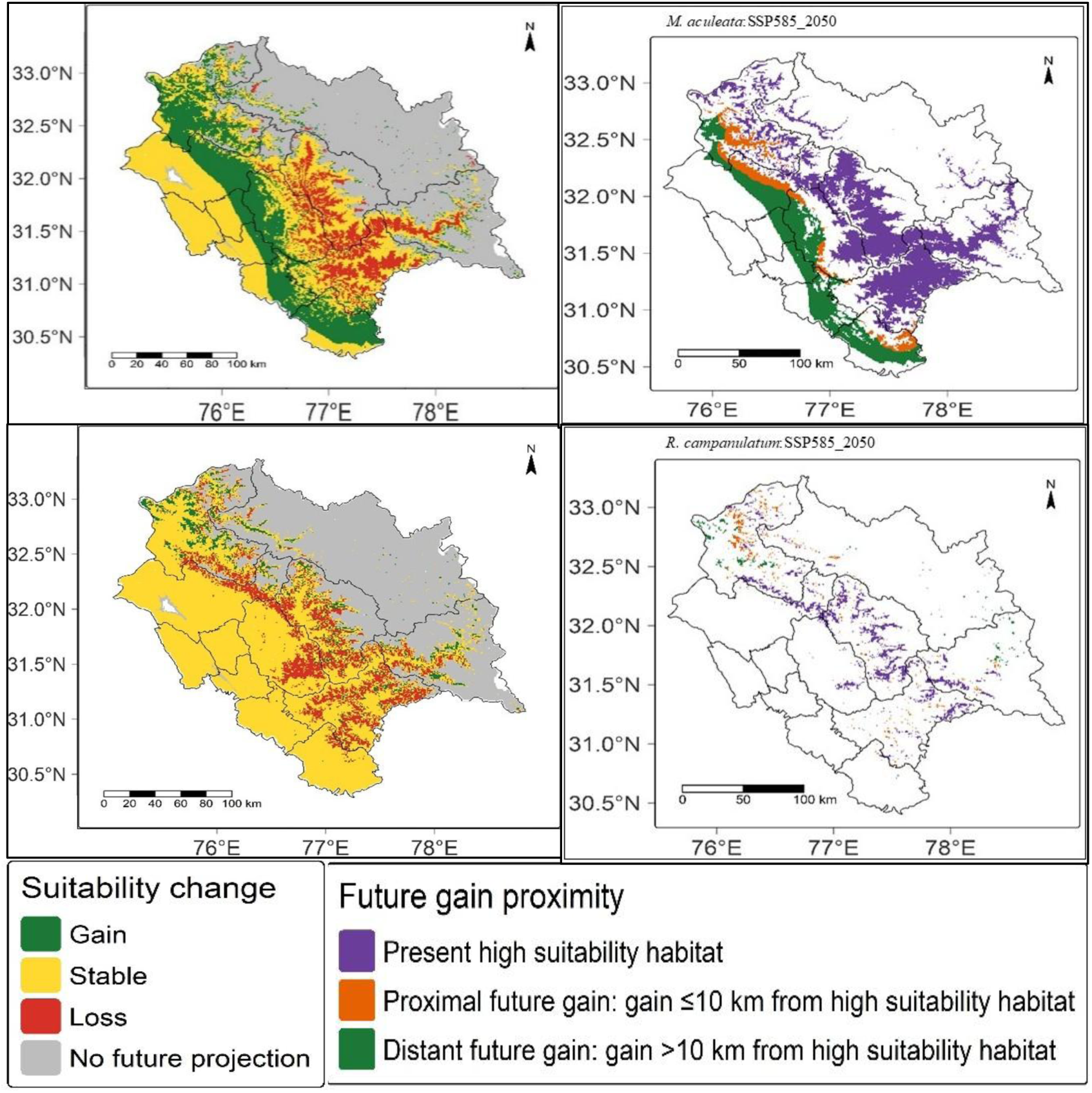
Suitability change (SSP 5-8.5, 2050) of *M. aculeata* (top right panel) and *R. campanulatum* (bottom right panel) and future grain proximity (SSP 5-8.5, 2050) of *M. aculeata* and *R. campanulatum*

To examine the spatial relationship between present high-suitability habitat and projected areas of future gain, gain areas were further classified according to their proximity to currently highly suitable habitat **(Online Resource 3, Fig. S9-S10)**. For *M. aculeata*, proximal future gain areas occurring within 10 km of present high-suitability habitat were concentrated primarily along the northwestern margin of the current distribution, with additional smaller areas towards its southern extent. However, a substantial proportion of projected gain occurred more than 10 km from currently highly suitable habitat, forming a relatively continuous band along the western and southwestern parts of the species’ distribution. This spatial pattern was broadly consistent across SSPs and future periods **(Fig. 5).**

In contrast, projected gain areas for *R. campanulatum* were comparatively limited and spatially fragmented. Gain areas located within 10 km of present high-suitability habitat occurred mainly around the northern and northwestern portions of the current distribution, with smaller scattered patches elsewhere. More distant gain areas were generally small and isolated. The spatial overlap of future proximal gain areas between the two species was very limited (**Online resource 4,** Fig. S11). Across the projected scenarios, areas classified as proximal gains for both species were negligible, with future proximal gains occurring almost exclusively in different parts of the study region. Overall, the proximity maps indicate contrasting spatial configurations of projected habitat gain between the two species: *M. aculeata* showed a mixture of proximal and more spatially separated future gain, whereas future gains for *R. campanulatum* were smaller and more fragmented, with proximal gain concentrated around particular portions of its existing high-suitability distribution.

### 3.5 Future range shift and centroid analysis

Centroid analyses indicated relatively consistent directional displacement of predicted suitable habitat distributions for both species under future climate scenarios.

For *M. aculeata*, the centroid of predicted suitable habitat shifted by 8.50 km towards the southwest (209.1°) under SSP1-2.6 and SSP2-4.5 for both 2050 and 2070. Under SSP5-8.5, centroid displacement was 7.74 km, with the direction of displacement being approximately southward (180.39°) in both 2050 and 2070. Schoener’s D values ranged from 0.507 to 0.674 across the projections. Values under SSP1-2.6 and SSP2-4.5 were approximately 0.673, whereas values under SSP5-8.5 were 0.507, indicating lower spatial overlap between current and projected suitable-habitat distributions under the highest-emission scenario.

For *R. campanulatum,* centroid displacement ranged from 8.41 to 8.50 km across the future scenarios and periods. The displacement was consistently directed towards the southwest, with bearings ranging from 209.12° to 209.84°. Schoener’s D values ranged from 0.687 to 0.689 across all scenarios and periods, indicating consistently high spatial overlap between current and projected suitable-habitat distributions.

Across both species, centroid displacement remained within approximately 8-9 km under all future scenarios. However, Schoener’s D showed greater variation among scenarios for *M. aculeata* than for *R. campanulatum*, particularly under SSP5-8.5, whereas the values for *R. campanulatum* remained within a narrow range across all projections **(Online Resource 2, Table S8)**.

## Discussion

The present study shows that future climate scenarios are projected to alter the distribution and availability of suitable habitat for *M. aculeata* and *R. campanulatum*. This spatial redistribution is structured by combinations of climatic, topographic, land-cover, and other environmental predictors. The MaxEnt model analyses identified elevation as the strongest individual predictor for both species, while several temperatures- and precipitation related climatic predictors were also important in both models. The response curves further showed that predicted suitability did not change uniformly across all environmental gradients. Together, these findings indicate that projected responses cannot be interpreted simply as direct responses to warming or precipitation change alone but instead reflect combinations of environmental relationships that differ in relative importance and in the shape of response patterns between the two species. Both species showed relatively small centroid displacements of approximately 8-9 km despite reductions in total suitable habitat under all future projections. Thus, the projected changes are characterized less by a large-scale displacement of the geographic centre of suitable habitat than by heterogeneous spatial redistribution, loss, persistence and gain of suitable areas across the landscape.

For *M. aculeata*, elevation provided the highest gain when used individually in the jackknife analysis, followed by precipitation seasonality (BIO15), mean temperature of the wettest quarter (BIO8), precipitation of the coldest quarter (BIO19), and mean diurnal temperature range (BIO2). Thus, both precipitation- and temperature-related variables were among the most important climatic predictors retained in the model. The full-model response curves indicate that these predictors were associated with predicted suitability in different ways. Suitability increased up to approximately 2,200 m and remained relatively stable until approximately 2,500 m before declining gradually at higher elevations. This response should not be interpreted as evidence that *M. aculeata* preferentially occupies lower elevation or subalpine habitats, as the observed field populations occupy areas between 3,264 and 4,176 m. Rather, the response curves represents the modelled relationships with elevation across the environmental space represented by the occurrence and background data and may also reflect correlations between elevation and other climatic predictors included in the model. The curve therefore describes the contribution of elevation within the fitted model rather than providing a direct ecological estimate of the species’ elevational optimum.

Among the climatic variables, *M. aculeata* showed responses to both precipitation- and temperature-related conditions. Predicted suitability declined with increasing precipitation seasonality, particularly above approximately 40, whereas suitability increased with precipitation during the coldest quarter up to approximately 220 mm before declining at higher values. At the same time, the response to mean diurnal temperature range declined with increasing values, while the response to mean temperature of the wettest quarter was relatively stable at lower temperatures before showing a pronounced peak at approximately 17 °C and declining at higher values. These results therefore indicate that the climatic component of the model reflects a combination of precipitation and temperature relationships rather than a predominantly temperature- or precipitation-driven response. This partly contrasts with generalized expectations that alpine species are necessarily limited primarily by temperature conditions and highlights the importance of considering the specific environmental relationships represented within individual species models (Körner, 2021). Previous work has likewise identified multiple environmental controls on the distribution of *M. aculeata* and we notice that the relative importance and direction of individual predictors differ among landscapes (Paul and Samant 2024b, the present study). We think that especially this aspect of landscape-dependence (of which environmental variables are the most important predictors for a given species) is very important for effective conservation planning and for the design of climate-proof protected areas. How the importance of different climatic variables depends on landscape needs to be more deeply understood and is thus deserving of systematic future investigation. Together with previous projections reporting shrinkage, expansion, or relatively stable suitability in different Himalayan landscapes, these differences suggest that the projected response of *M. aculeata* cannot be reduced to a single species-wide response to climate change. Rather, the outcome appears to depend on the environmental conditions represented within the particular landscape being modelled.

For *R. campanulatum*, elevation likewise provided the highest individual gain in the jackknife analysis followed closely by mean diurnal temperature range (BIO2). Land cover, and several precipitation- and temperature-related variables, including BIO15, BIO19 and BIO8, also contributed to the model. The response curves similarly indicate a strong association with elevation, with predicted suitability increasing up to approximately 3,200 m, and declining sharply at higher elevations. This response is broadly consistent with the observed occurrence of the species within the subalpine elevational zone, although the field populations covered a relatively broad elevational range.

The climatic response of *R. campanulatum* also involved both temperature- and precipitation-related variables. Predicted suitability declined strongly with increasing mean diurnal temperature range, particularly between approximately 12 and 14 °C. Suitability increased with isothermality up to approximately 33.5 before declining at higher values. The response to mean temperature of the wettest quarter showed a more complex pattern, with two peaks at approximately -7.5 and 14 °C rather than a single narrow optimum. At the same time, suitability declined with increasing precipitation seasonality, while the response to precipitation during the coldest quarter peaked around 220 mm before declining. These response patterns indicate that the climatic component of the *R. campanulatum* model was also associated with a combination of temperature variability and precipitation conditions rather than with a purely temperature-driven response.

The comparison between the two species therefore indicates substantial overlap in the environmental dimensions represented by their models. Elevation was the strongest individual predictor for both species, and several climatic variables, including BIO2, BIO15 and BIO19, were important for both models, although their relative contributions differed. The response curves also showed broadly comparable non-linear relationships for several predictors, including declining suitability at higher BIO15 values and unimodal responses to BIO19. The main differences therefore lie less in a clear precipitation versus temperature separation than in the relative importance and specific shapes of individual predictor responses. For example, BIO2 showed a particularly strong association with *R. campanulatum*, whereas *M. aculeata* showed comparatively stronger associations with BIO15, BIO8, and BIO19 relative to its other predictors. These differences indicate that similar projected reductions in total suitable area do not necessarily arise from identical environmental relationships and responses.

The field surveys provide additional ecological context for interpreting these modelled relationships. *M. aculeata* occurred in small and localized populations within open alpine herbaceous communities, whereas *R. campanulatum* formed extensive stands within subalpine shrub communities. The two habitats also differed substantially in associated species composition, with only 14 associated plant species shared between the two communities. The field data therefore demonstrate that the two focal species occupy rather distinct vegetation contexts across the alpine-subalpine transition. These observations complement the distribution models by describing population occurrence, abundance, frequency and associated plant communities; information that is not provided by habitat-suitability modelling alone.

The contrasting community associations may be relevant to future redistribution. *M. aculeata* occurs within open alpine herbaceous communities, whereas *R. campanulatum* is associated with shrub-dominated subalpine vegetation. Future changes in habitat suitability therefore occur within ecological systems that may themselves change in vegetation structure and composition. For example, shifts in the distribution of woody vegetation, changes in associated species, or alterations in land use and grazing pressure could modify the conditions under which suitable climatic and topographic environments are occupied. These processes are not represented directly in the present models and require long-term ecological monitoring in the field.

Both species showed substantial reductions in total suitable habitat under future climate scenarios, although the magnitude of change was relatively similar across the three emissions pathways. For *M. aculeata*, suitable habitat declined from 2,785.82 km² under current conditions to approximately 2,076-2,090 km² across the future projections, corresponding to reductions of approximately 25%. For *R. campanulatum*, suitable habitat declined from 777.89 km² to approximately 558-565 km², corresponding to reductions of approximately 27-28%. The relatively small differences among scenarios suggest that, within the projected environmental space represented by the HadGEM3-GC31-LL projections and the selected predictors, the overall area of suitable habitat changed to a broadly similar extent across the three SSPs.

Importantly, the suitability-change maps show that these reductions do not represent uniform habitat loss throughout the landscape. Both species exhibited a mosaic of projected loss, persistence, and gain. For *M. aculeata*, future gain areas formed a relatively broad band, particularly towards the western and southwestern parts of the distribution, while *R. campanulatum* showed smaller and more spatially concentrated areas of projected gain. These results imply that projected habitat redistribution is both species-specific and landscape-dependent rather than representing a simple, uniform upslope movement.

The proximity analysis provides additional information about the spatial configuration of these projected gain areas. For *M. aculeata*, some future gain areas occurred within 10 km of currently highly suitable habitat, particularly along the northwestern margin of the distribution and in smaller areas towards its southern extent. However, a substantial proportion of projected gain was located farther than 10 km from currently highly suitable areas, forming a more continuous band across the western and southwestern parts of the distribution. In contrast, projected gain for *R. campanulatum* was comparatively limited and fragmented, with proximal gains occurring mainly around the northern and northwestern portions of the existing distribution.

These spatial patterns are relevant because projected gain does not necessarily imply that newly suitable areas will be colonized. Areas located closer to existing highly suitable habitat may potentially be more accessible to natural range adjustment than spatially separated areas, although actual colonization will depend on dispersal, recruitment, intervening landscape conditions, and biotic interactions. The very limited overlap between future proximal gain areas of the two species also has implications for spatial conservation planning. Previous multi-taxa conservation-planning research has shown that spatial priorities can differ among taxa and that incorporating projected future distributions can alter the representation of biodiversity within protected-area networks (Critchlow et al. 2022). If conservation planning aims to protect both focal species and their associated habitats, the spatial separation of their projected proximal gains suggests that a single conservation area may not adequately encompass future suitable areas for both species. Conservation strategies may therefore need to consider a network of complementary areas or, where feasible, larger protected areas that encompass the distinct locations of future proximal habitat gains. The present analysis does not quantify dispersal or functional connectivity, and the 10-km threshold should therefore be interpreted as a spatial proximity measure rather than as evidence that a species can disperse across this distance. Nevertheless, distinguishing between proximal and more spatially separated future gain areas provides a more informative picture than considering total habitat gain alone and identifies locations that could be prioritized to enhance connectivity within and between protected areas. The centroid results further demonstrate that changes in the spatial configuration of suitable habitat can occur without large displacement of its geographic centre (Casazza et al. 2023) . For both species, centroid displacement remained within approximately 8-9 km across future scenarios. However, Schoener’s D showed greater variation for *M. aculeata* than for *R. campanulatum*. For *M. aculeata*, D was approximately 0.67 under SSP1-2.6 and SSP2-4.5 but declined to approximately 0.51 under SSP5-8.5, whereas *R. campanulatum* maintained values of approximately 0.69 across all scenarios. In the present analysis, these values represent spatial overlap between current and projected habitat-suitability distributions rather than a direct measure of climatic niche breadth or stability. The lower overlap for *M. aculeata* under SSP5-8.5 therefore indicates greater spatial reorganization of predicted suitable habitat under that scenario but does not by itself demonstrate that the species is intrinsically more sensitive to climate change than *R. campanulatum*.

The combination of habitat contraction, spatial redistribution, and relatively small centroid displacement has implications for conservation planning. The results suggest that future change may involve the progressive loss and reorganization of suitable habitat rather than a simple relocation of the entire distribution towards a new geographic centre. Conservation assessments should therefore consider not only where high suitability occurs under current or future conditions separately, but also the spatial relationship between present high-suitability areas and projected future gains. Areas where current suitable habitat persists and where future suitable or gain areas occur nearby may represent particularly relevant candidates for further assessment because they could provide greater opportunities for population persistence or natural range adjustment.

However, these areas should not automatically be described as climate refugia. The present analysis identifies locations where suitable habitat persists under future projections or where future suitability is gained, but it does not directly demonstrate climatic or microclimatic buffering. Similarly, the ecological value of candidate areas may differ between the two species because their models respond to different combinations of environmental predictors and their projected gain areas have different spatial configurations. The results also raise questions about the use of individual species as ecological indicators. Species are likely to differ in the relative importance of different predictors and in their response curves for these predictors. This suggests that the response of one species cannot automatically be assumed to represent, as an indicator, the response of another species or of the wider alpine flora. The present analysis does not formally quantify the effectiveness of the two species as indicators for other spcies, and therefore does not establish that either species can serve as a general indicator for Himalayan alpine biodiversity. Rather, the results demonstrate that examining ecologically contrasting species can reveal different spatial and environmental dimensions of projected habitat change, including how changes in both temperature- and precipitation-related variables interact with elevation to affect habitat suitability for alpine and subalpine plant species.

Future work could build on this comparison by explicitly testing how conservation recommendations differ when they are based on individual species versus combinations of species with contrasting environmental associations. Such analyses could quantify the extent to which conservation priorities derived from one species overlap with, or fail to represent, suitable habitat for others. Extending this approach to additional alpine and subalpine species would provide a stronger basis for evaluating whether complementary sets of species offer more informative indicators of landscape-level environmental change than individual species considered in isolation.

Conservation prioritization would benefit from incorporating additional species, population trends, functional and community-level information, habitat connectivity, and processes such as land use, grazing, harvesting and vegetation change that are not represented directly in the present models.

Overall, the principal finding of this study is that projected habitat change in the Western Himalaya is spatially heterogeneous and species-specific. *M. aculeata* and *R. campanulatum* show similar overall reductions in suitable habitat and share several important climatic predictors but differ in the relative importance and response patterns of individual environmental variables, spatial overlap between present and future suitability, and configuration of projected habitat gains. Our findings emphasize that projected climate responses may not always occur as simple directional range shifts. Instead, species-specific environmental associations and landscape structure appear to be important for how suitable habitat may be redistributed under climate change.

## Conclusion

This study combined field-based population and habitat observations with species distribution modelling to assess current habitat associations and projected changes in suitable habitat for *M. aculeata* and *R. campanulatum* in Himachal Pradesh. Field surveys showed contrasting ecological and population settings for the two species: *M. aculeata* occurred in small, localized populations within open alpine herbaceous communities, whereas *R. campanulatum* occurred across more widespread subalpine shrub communities. These field observations provide ecological context for interpreting the spatial predictions of the species distribution models.

The MaxEnt analyses indicated that habitat suitability was strongly associated with elevation and multiple climatic predictors for both species. Several important temperature- and precipitation-related predictors were shared between the two models, including BIO2, BIO15 and BIO19, demonstrating that the environmental relationships of the species cannot be separated into a simple temperature-driven versus precipitation-driven contrast. Instead, the species differed primarily in the relative importance and response patterns of individual predictors. *R. campanulatum* showed particularly strong associations with elevation and mean diurnal temperature range, whereas *M. aculeata* showed relatively stronger associations with precipitation seasonality, mean temperature of the wettest quarter, and precipitation of the coldest quarter. Both models nevertheless indicate that predicted habitat suitability reflects combinations of temperature, precipitation, topographic and other environmental conditions rather than a single dominant climatic control.

Future projections indicated a reduction in total suitable habitat for both species across SSP1- 2.6, SSP2-4.5 and SSP5-8.5. Suitable habitat for *M. aculeata* declined by approximately 25% relative to current conditions, whereas the decline for *R. campanulatum* was approximately 27-28% by 2050-2070. The habitat-change maps further showed that these reductions comprise spatially heterogeneous areas of habitat loss, gain and persistence rather than uniform contraction across Himachal Pradesh. Centroid displacement remained relatively small, at approximately 8-9 km, while Schoener’s D indicated greater changes in the spatial overlap of current and future suitable habitat for *M. aculeata* under SSP5-8.5 than under the lower-emission scenarios. In contrast, Schoener’s D remained relatively stable for *R. campanulatum* across scenarios.

Together, these findings indicate that climate-related change in habitat suitability for the two species is likely to occur primarily through gradual and spatially heterogeneous changes in habitat availability and configuration rather than through large displacement of the geographic centre of suitable habitat. Although the two species share several important environmental predictors and experience broadly similar overall reductions in suitable habitat, their individual predictor relationships and projected spatial configurations differ.

From a conservation perspective, the spatial relationship between current high-suitability habitat and projected areas of future gain may help identify locations where range adjustment by natural dispersal is more feasible. The contrasting configurations of proximal and more distant future gain observed for the two species further indicate that the accessibility of newly suitable habitat may differ between species. Areas where suitable habitat persists, or where projected future gain occurs close to currently highly suitable habitat, represent candidate areas for further conservation assessment. Long-term field monitoring and maintenance of habitat connectivity should complement predictive modelling. Conservation planning should also incorporate additional alpine and subalpine species before broader biodiversity-level priorities are established.

## Statements and Declarations Funding

The authors declare that no funds, grants, or other support were received during the preparation of this manuscript.

## Declaration of competing interest

The authors declare that they have no known competing financial interests or personal relationships that could have appeared to influence the work reported in this paper.

## Author contributions

Material preparation, data collection, and analysis were performed by **Simran Tomar**. The first draft of the manuscript was written by **Simran Tomar**. **Matthijs Vos** contributed to the conceptualization of the analyses and reviewed and edited the initial drafts of the manuscript. **Merja Helena Tolle** reviewed and edited subsequent drafts of the manuscript.

## Data availability

The author confirms that all data generated or analyzed during this study are included in this published article. Furthermore, primary and secondary sources and data supporting the findings of this study were all publicly available at the time of submission. Any additional datasets generated during and/or analysed during the current study are available from the corresponding author on reasonable request.

## Notes

### Competing Interest Statement

The authors have declared no competing interest.

## References

1. Ahmad M, Luo Y-H, Rathee S, Spicer RA, Zhang J, Wambulwa MC, Zhu G-F, Cadotte MW, Wu Z-Y, Khan SM, Maity D, Li D-Z, Liu J (2025) Multifaceted plant diversity patterns across the Himalaya: Status and outlook. Plant Divers. 10.1016/j.pld.2025.04.003

2. Allouche O, Tsoar A, Kadmon R (2006) Assessing the accuracy of species distribution models: Prevalence, kappa and the true skill statistic (TSS). Journal of Applied Ecology 43:1223–1232. 10.1111/j.1365-2664.2006.01214.x

3. Basnett S, Ganesan R (2022) A Comprehensive Review on the Taxonomy, Ecology, Reproductive Biology, Economic Importance and Conservation Status of Indian Himalayan Rhododendrons. The Botanical Review 88:505–544. 10.1007/s12229-021-09273-z

4. Booth TH (2022) Checking bioclimatic variables that combine temperature and precipitation data before their use in species distribution models. Austral Ecol 47:1506–1514. 10.1111/aec.13234

5. Casazza G, Guerrina M, Dagnino D, Minuto L (2023) Will natura 2000 european network of protected areas support conservation of Southwestern Alps endemic flora under future climate? Biodivers Conserv 32:1353–1367. 10.1007/s10531-023-02556-4

6. Chauhan S, Ghoshal S, Kanwal KS, Sharma V, Ravikanth G (2022) Ecological niche modelling for predicting the habitat suitability of endangered tree species Taxus contorta Griff. in Himachal Pradesh (Western Himalayas, India). Trop Ecol 63:300–313. 10.1007/s42965-021-00200-2

7. Critchlow R, Cunningham CA, Crick HQP, Macgregor NA, Morecroft MD, Pearce-Higgins JW, Oliver TH, Carroll MJ, Beale CM (2022) Multi-taxa spatial conservation planning reveals similar priorities between taxa and improved protected area representation with climate change. Biodivers Conserv 31:683–702. 10.1007/s10531-022-02357-1

8. Elith J, Leathwick JR (2009) Species distribution models: Ecological explanation and prediction across space and time. Annu Rev Ecol Evol Syst 40:677–697. 10.1146/annurev.ecolsys.110308.120159

9. Fick SE, Hijmans RJ (2017) WorldClim 2: new 1-km spatial resolution climate surfaces for global land areas. International Journal of Climatology 37:4302–4315. 10.1002/joc.5086

10. Franklin J (2010) Mapping Species Distributions. Cambridge University Press

11. Ganaie H, Ahmad D, Kaloo Z, Ganai B, Singh S (2016) Phytochemical Screening of Meconopsis aculeata Royle an Important Medicinal Plant of Kashmir Himalaya: A Perspective. Research Journal of Phytochemistry 10. 10.3923/rjphyto.2016.

12. GBIF Occurrence (2023) GBIF.org. https://www.gbif.org/occurrence/search?q=ACONITUM%20HETEROPHYLLUM&continent=ASIA&country=IN. Accessed 27 Apr 2024

13. Hooker JD (1890) The flora of british India Johnson B, Pinilla-Buitrago GE, Paz A Wallace Ecological Modelling App. In: 2022

14. Kala CP (2003) Medicinal plants of Indian trans-Himalaya: focus on Tibetan use of medicinal resources. Bishen Singh Mahendra Pal Singh

15. Kattel GR (2022) Climate warming in the Himalayas threatens biodiversity, ecosystem functioning and ecosystem services in the 21st century: is there a better solution? Biodivers Conserv 31:2017–2044. 10.1007/s10531-022-02417-6

16. Körner C (2003) Alpine Plant Life. Springer Berlin Heidelberg, Berlin, Heidelberg

17. Kumar P, Sarthi PP (2021) Intraseasonal variability of Indian Summer Monsoon Rainfall in CMIP6 models simulation. Theor Appl Climatol 145:687–702. 10.1007/s00704-021-03661-6

18. Kumar S, Khanduri VP (2024) Impact of climate change on the Himalayan alpine treeline vegetation. Heliyon 10:e40797. 10.1016/j.heliyon.2024.e40797

19. Majid A, Ahmad H, Saqib Z, Ali H, Alam J (2015) Conservation status assessment of Meconopsis aculeata Royle: A threatened endemic of Pakistan and Kashmir. Pak J Bot 47:1– 5

20. Manzoor M, Ahmad M, Gillani SW, Waheed M, Shaheen H, Mehmood AB, Fonge BA, Al-Andal A (2025) Population dynamics, threat assessment, and conservation strategies for critically endangered Meconopsis aculeata in alpine zone. BMC Plant Biol 25:358. 10.1186/s12870-025-06361-9

21. Meher JK, Das L (2024) Probabilistic evaluation of three generations of climate models for simulating precipitation over the Western Himalayas. Journal of Earth System Science 133:15. 10.1007/s12040-023-02216-9

22. Merow C, Smith MJ, Silander JA (2013) A practical guide to MaxEnt for modeling species’ distributions: what it does, and why inputs and settings matter. Ecography 36:1058–1069. 10.1111/j.1600-0587.2013.07872.x

23. Misra R (1968) Ecology Work Book. Oxford and IBH Publishing Co., Calcutta

24. Myers N, Mittermeier RA, Mittermeier CG, da Fonseca GAB, Kent J (2000) Biodiversity hotspots for conservation priorities. Nature 403:853–858. 10.1038/35002501

25. Ohlemüller R, Anderson BJ, Araújo MB, Butchart SHM, Kudrna O, Ridgely RS, Thomas CD (2008) The coincidence of climatic and species rarity: high risk to small-range species from climate change. Biol Lett 4:568–572. 10.1098/rsbl.2008.0097

26. O’Neill BC, Kriegler E, Ebi KL, Kemp-Benedict E, Riahi K, Rothman DS, van Ruijven BJ, van Vuuren DP, Birkmann J, Kok K, Levy M, Solecki W (2017) The roads ahead: Narratives for shared socioeconomic pathways describing world futures in the 21st century. Global Environmental Change 42:169–180. 10.1016/j.gloenvcha.2015.01.004

27. Paul S, Samant SS (2024a) Population ecology and habitat suitability modelling of an endangered and endemic medicinal plant Meconopsis aculeata Royle under projected climate change in the Himalaya. Environ Exp Bot 225:105837. 10.1016/j.envexpbot.2024.105837

28. Paul S, Samant SS (2024b) Population ecology and habitat suitability modelling of an endangered and endemic medicinal plant Meconopsis aculeata Royle under projected climate change in the Himalaya. Environ Exp Bot 225:105837. 10.1016/j.envexpbot.2024.105837

29. Pepin NC, Arnone E, Gobiet A, Haslinger K, Kotlarski S, Notarnicola C, Palazzi E, Seibert P, Serafin S, Schöner W, Terzago S, Thornton JM, Vuille M, Adler C (2022) Climate Changes and Their Elevational Patterns in the Mountains of the World. Reviews of Geophysics 60. 10.1029/2020RG000730

30. Phillips SJ, Anderson RP, Schapire RE (2006) Maximum entropy modeling of species geographic distributions. Ecol Modell 190:231–259. 10.1016/j.ecolmodel.2005.03.026

31. Sekar KC, Thapliyal N, Bhojak P, Bisht K, Pandey A, Mehta P, Negi VS, Rawat RS (2025) Early signals of climate change impacts on alpine plant diversity in Indian Himalaya. Biodivers Conserv 34:207–233. 10.1007/s10531-024-02966-y

32. Shaheen H, Aziz S, Nasar S, Waheed M, Manzoor M, Siddiqui MH, Alamri S, Haq SM, Bussmann RW (2023) Distribution patterns of alpine flora for long-term monitoring of global change along a wide elevational gradient in the Western Himalayas. Glob Ecol Conserv 48:e02702. 10.1016/j.gecco.2023.e02702

33. Shukla V, Singh A, Nautiyal AR (2021a) Population Assessment and Phyto-chemical Screening of Meconopsis aculeata Royle an Endangered Medicinal Plant of Western Himalaya. In: Springer Proceedings in Earth and Environmental Sciences. Springer Nature, pp 429–443

34. Shukla V, Singh A, Nautiyal AR (2021b) Population Assessment and Phyto-chemical Screening of Meconopsis aculeata Royle an Endangered Medicinal Plant of Western Himalaya. In: Springer Proceedings in Earth and Environmental Sciences. Springer Nature, pp 429–443

35. Singh N, Tewari A, Shah S, Mittal A, Gangola S, Wani ZA (2025) Seasonal dynamics and adaptation strategies of krummholz forming Rhododendron campanulatum to water availability at high-altitude Himalayan treeline environments. PLoS One 20:e0318197. 10.1371/journal.pone.0318197

36. Steven J. Phillips, Miroslav Dudík, Robert E. Schapire (2022) Maxent software for modeling species niches and distributions

37. Swets JA (1988) Measuring the Accuracy of Diagnostic Systems. Science (1979) 240:1285–1293. 10.1126/science.3287615

38. Tiwary R, Singh PP, Adhikari D, Behera MD, Barik SK (2024) Vulnerability assessment of Taxus wallichiana in the Indian Himalayan Region to future climate change using species niche models and global climate models under future climate scenarios. Biodivers Conserv 33:3475–3494. 10.1007/s10531-024-02859-0

39. Warren DL, Richard EG, Michael T (2008) ENVIRONMENTAL NICHE EQUIVALENCY VERSUS CONSERVATISM: QUANTITATIVE APPROACHES TO NICHE EVOLUTION. Evolution (N Y) 62:1215–1215. 10.1111/j.1558-5646.2010.01204.x

