## Supplementary files_1 for "Projected habitat loss and spatial redistribution of two alpine and subalpine species in Himachal Pradesh, Western Himalaya under CMIP6 scenarios": ESM_1.pdf

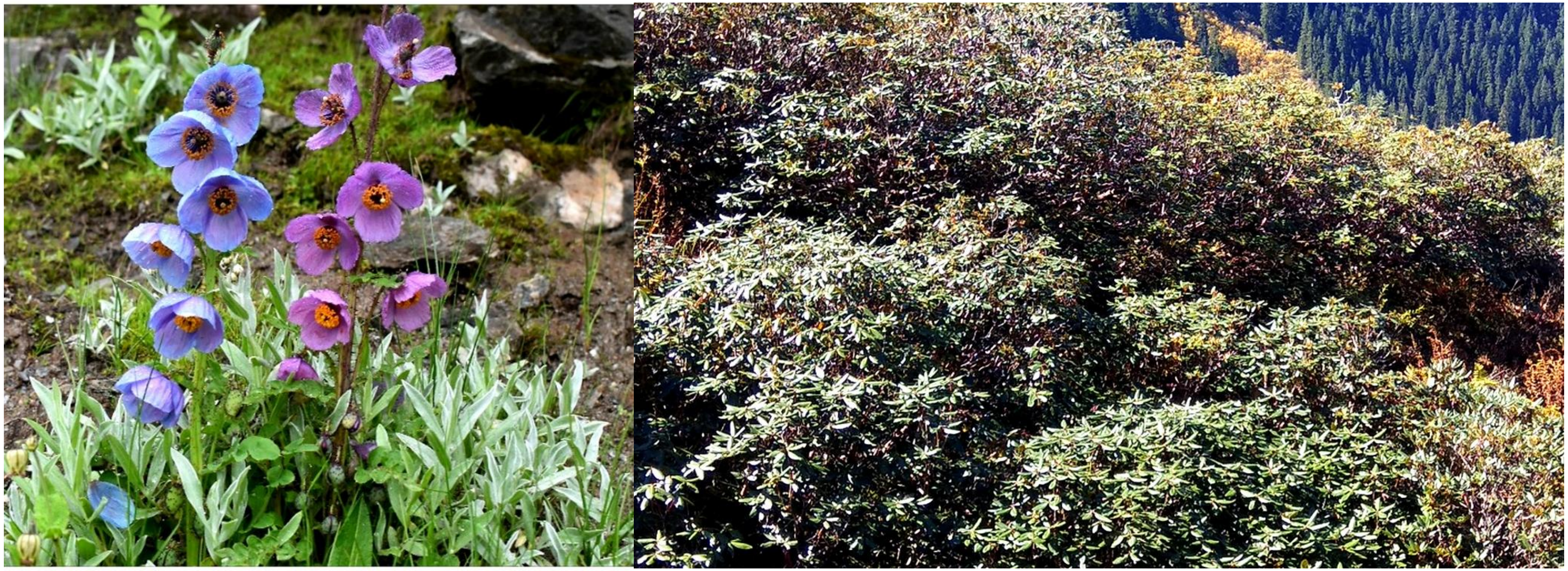

**Fig. S1** *M. aculeata* (*left panel*) and R. campanulatum (*right panel*) in their natural habitat near Rohtang Pass, Beas valley and Parvati valley

**Table S1.** Ethnomedicinal and traditional uses of *M. aculeata* and *R. campanulatum.*

| Species | Plant Part Used | Ethnomedicinal / Traditional Use | Bio-active Compounds | References |
|---|---|---|---|---|
| *M. aculeata* | Flowers | Floral paste for body aches | Polyphenols, flavonoids, terpenoids, alkaloids, phytosterols | (Bahukhandi et al., 2019; Ganaie et al., 2016; Shukla et al., 2022) |
| | Leaves | Leaf pastes for swollen legs | - | (Hadi and Singh, 2021) |
| | Whole plant | Powder to enhance immunity during childbirth; treatment of rheumatic pain, fever, asthma, kidney ailments, infections | - | (Jan et al., 2023; Kala, 2003) (Aswal and Mehrotra, 1994; Tsarong and Tsewang, 1994) |
| *R. campanulatum* | Leaves | Treatment of syphilis, rheumatism, skin diseases | Phenolics, flavonoids, tannins | 2013; (Bhattacharyya, 2011; Kala et al., 2004; Popescu and Kopp, 2013) |
| | Twigs & Wood | Brewed with tea for persistent fevers; fuelwood | - | Popescu & Kopp, 2013 |

**Table S2**. Site characteristics of the selected populations of *M. aculeata* and *R. campanulatum*

| **Species** | **Site** | **Elevation (m asl)** | **Slope (°)** | **Aspect** | **Habitat** |
|---|---|---|---|---|---|
| ***M. aculeata*** | Rohtang Pass | 3264 | 65 | NW | Bouldery |
| | Near Rani Nala, Rohtang Pass | 4176 | 60 | S | Alpine grassland |
| ***R. campanulatum*** | Fozal, Beas valley | 2850 | 65 | NW | Bouldery |
| | Fozal, Beas valley | 3136 | 50 | N | Shady moist |
| | Fozal, Beas valley | 3209 | 60 | N | Dry |
| | Beas valley | 3851 | 65 | NW | Dry |
| | Parvati valley | 3306 | 65 | NW | Shady moist |
| | Parvati valley | 3360 | 50 | SE | Bouldery |
| | Parvati valley | 2934 | 50 | NE | Shady moist |
| | Parvati valley | 3523 | 45 | N | Shady moist |
| | Parvati valley | 3624 | 35 | N | Dry |
| | Jalori pass, Banjar | 3227 | 77 | SE | Open forest |
| | Near Lambri top | 3200 | 57 | S | Open forest |
| | Lambri top, Sarchi | 3500 | 73 | S | Rock scree |

**Table S3. Species occurrence record and their sources.**

| S.no | Species | Latitude | Longitude | Source |
|---|---|---|---|---|
| ***R. campanulatum*** | | | | |
| 1. | *R. campanulatum* | 31.34095 | 78.46092 | GBIF |
| 2. | *R. campanulatum* | 32.08496 | 77.05154 | Field based |
| 3. | *R. campanulatum* | 32.04546 | 77.04437 | GBIF |
| 4. | *R. campanulatum* | 32.04641 | 77.04307 | GBIF |
| 5. | *R. campanulatum* | 31.27472 | 77.13863 | GBIF |
| 6. | *R. campanulatum* | 31.98403 | 77.50257 | Field based |
| 7. | *R. campanulatum* | 31.99061 | 77.50777 | Field based |
| 8. | *R. campanulatum* | 31.97772 | 77.48762 | Field based |
| 9. | *R. campanulatum* | 31.97637 | 77.48162 | Field based |
| 10. | *R. campanulatum* | 30.37760 | 77.28132 | GBIF |
| 11. | *R. campanulatum* | 30.37764 | 77.28192 | GBIF |
| 12. | *R. campanulatum* | 30.37683 | 77.28203 | GBIF |
| 13. | *R. campanulatum* | 30.37716 | 77.28132 | GBIF |
| 14. | *R. campanulatum* | 30.37737 | 77.28154 | GBIF |
| 15. | *R. campanulatum* | 30.37739 | 77.28158 | GBIF |
| 16. | *R. campanulatum* | 32.08033 | 77.05357 | Field based |

| | | | | |
|---|---|---|---|---|
| 17. | *R. campanulatum* | 32.04546 | 77.04437 | Field based |
| 18. | *R. campanulatum* | 32.04641 | 77.04307 | GBIF |
| 19. | *R. campanulatum* | 32.37767 | 77.28135 | Field based |
| 20. | *R. campanulatum* | 32.32506 | 76.30767 | GBIF |
| 21. | *R. campanulatum* | 32.31954 | 76.30701 | GBIF |
| 22. | *R. campanulatum* | 32.16635 | 72.76375 | GBIF |
| 23. | *R. campanulatum* | 32.08421 | 76.92452 | GBIF |
| 24. | *R. campanulatum* | 32.08421 | 76.92452 | GBIF |
| 25. | *R. campanulatum* | 31.20254 | 77.91884 | Field based |
| 26. | *R. campanulatum* | 31.18992 | 78.10681 | GBIF |
| 27. | *R. campanulatum* | 31.19803 | 78.11732 | GBIF |
| 28. | *R. campanulatum* | 31.591062 | 77.402022 | GBIF |
| 29. | *R. campanulatum* | 31.595292 | 77.595292 | Field based |
| 30. | *R. campanulatum* | 31.539951 | 77.38751 | Field based |
| 31. | *R. campanulatum* | 34.08 | 74.82 | GBIF |
| 32. | *R. campanulatum* | 33.0000000 | 77.0000000 | GBIF |
| 33. | *R. campanulatum* | 34.030 | 74.350000 | GBIF |
| 34. | *R. campanulatum* | 30.486202 | 79.217722 | GBIF |
| 35. | *R. campanulatum* | 30.701621 | 77.8696 | Field based |

| | | | | |
|---|---|---|---|---|
| 36. | *R. campanulatum* | 34.152442 | 74.981722 | GBIF |
| 37. | *R. campanulatum* | 30.063636 | 80.203011 | GBIF |
| 38. | *R. campanulatum* | 29.847342 | 80.536906 | GBIF |
| 39. | *R. campanulatum* | 31.340946 | 78.460922 | GBIF |
| 40. | *R. campanulatum* | 22.88361 | 79.61611 | GBIF |
| 41. | *R. campanulatum* | 22.883478 | 79.616202 | GBIF |
| ***M. aculeata*** | | | | |
| 1. | *M. aculeata* | 32.571 | 77.03198 | GBIF |
| 2. | *M. aculeata* | 31.50224 | 77.30607 | GBIF |
| 3. | *M. aculeata* | 31.59116 | 78.27659 | GBIF |
| 4. | *M. aculeata* | 32.8242 | 77.4477 | GBIF |
| 5. | *M. aculeata* | 32.05684 | 77.25514 | GBIF |
| 6. | *M. aculeata* | 31.80429 | 76.98831 | GBIF |
| 7. | *M. aculeata* | 31.69812 | 77.41623 | GBIF |
| 8. | *M. aculeata* | 31.41639 | 78.26972 | GBIF |
| 9. | *M. aculeata* | 31.55556 | 77.28556 | GBIF |
| 10. | *M. aculeata* | 31.415 | 77.2575 | GBIF |
| 11. | *M. aculeata* | 31.698124 | 77.416228 | GBIF |
| 12. | *M. aculeata* | 31.368629 | 78.136086 | GBIF |

| | | | | |
|---|---|---|---|---|
| 13. | *M. aculeata* | 31.183333 | 77.633333 | GBIF |
| 14. | *M. aculeata* | 30.32 | 80.18 | GBIF |
| 15. | *M. aculeata* | 30.733333 | 79.05 | GBIF |
| 16. | *M. aculeata* | 34.57 | 73.5 | GBIF |
| 17. | *M. aculeata* | 32.35 | 77.23 | Field based |
| 18. | *M. aculeata* | 32.37 | 77.25 | Field based |
| 19. | *M. aculeata* | 31.383333 | 78.5 | GBIF |
| 20. | *M. aculeata* | 33.9784 | 77.85294 | GBIF |
| 21. | *M. aculeata* | 34.87 | 75.12 | GBIF |

**Table S4.** List of variables used to generate Species Distribution Model.

| **Environmental variable** | **Resolution** | **Source** |
|---|---|---|
| **Bioclimatic variables** | | |
| Mean Diurnal Range (BIO2) | 30 arc-seconds ~1 kilometre | WorldClim; https://www.worldclim.org/data/BIOclim.html |
| Isothermality (BIO3=(BIO2/ BIO7) ×100) where BIO7 is temperature annual range | | |

| | | |
|---|---|---|
| Mean Temperature of Wettest Quarter (BIO8) | | |
| Precipitation Seasonality (Coefficient of Variation) (BIO15) | | |
| Precipitation of Coldest Quarter (BIO19), | | |
| **Topographic variable** | | |
| Elevation (Meters Above Sea Level) | 30 arc-seconds ~1 kilometre | (derived from SRTM elevation data) https://www.worldclim.org/data/worldclim21.html |
| Slope | 30 arc-seconds | Derived from elevation |
| Aspect | 30 arc-seconds | Derived from elevation |
| Soil Cover | 1 km resolution | Harmonized World Soil Database (HWSD) |
| Land Cover | 300 meters | ESA CCI Land Cover data, processed in R ('geodata' package) |

**Table S5** Environmental variables initially considered for species distribution modelling and their retention status following correlation and multicollinearity screening. Variables with pairwise Spearman correlation coefficients exceeding |r| = 0.70 were removed. Remaining predictors were evaluated using variance inflation factor (VIF) analysis, and all retained variables exhibited VIF values below 4.

| **Initial set of 24 variables** | **Variable removed/retained after correlation analysis** | **VIF** |
|---|---|---|
| BIO1 - Annual Mean Temperature | Removed | VIF values >10 |
| BIO2 - Mean Diurnal Range (Mean of monthly (max temp - min temp)) | Retained | 2.27 |
| BIO3 - Isothermality (BIO2/BIO7) (×100) | Retained | 3.58 |
| BIO4 - Temperature Seasonality (standard deviation ×100) | Removed | VIF values >10 |
| BIO5 - Max Temperature of Warmest Month | Removed | VIF values >10 |
| BIO6 - Min Temperature of Coldest Month | Removed | VIF values >10 |
| BIO7 - Temperature Annual Range (BIO5-BIO6) | Removed | VIF values >10 |
| BIO8 - Mean Temperature of Wettest Quarter | Retained | 2.15 |
| BIO9 - Mean Temperature of Driest Quarter | Removed | VIF values >10 |
| BIO10 - Mean Temperature of Warmest Quarter | Removed | VIF values >10 |
| BIO11 - Mean Temperature of Coldest Quarter | Removed | VIF values >10 |
| BIO12 - Annual Precipitation | Removed | VIF values >10 |

| | | |
|---|---|---|
| BIO13 - Precipitation of Wettest Month | Removed | VIF values >10 |
| BIO14 - Precipitation of Driest Month | Removed | VIF values >10 |
| BIO15 - Precipitation Seasonality (Coefficient of Variation) | Retained | 2.72 |
| BIO16 - Precipitation of Wettest Quarter | Removed | VIF values >10 |
| BIO17- Precipitation of Driest Quarter | Removed | VIF values >10 |
| BIO18 - Precipitation of Warmest Quarter | Removed | VIF values >10 |
| BIO19 - Precipitation of Coldest Quarter | Retained | 3.33 |
| Elevation | Retained | 3.23 |
| Slope | Retained | 1.70 |
| Aspect | Retained | 1.00 |
| Landcover | Retained | 1.83 |
| Soil | Retained | 1.36 |

**Table S6.** Phytosociological characteristics of *M. aculeata* and *R. campanulatum* populations in Himachal Pradesh

| Species | Location | Density [1] | Diversity (H') | Frequency, (F) (%) | Abundance, (A) | A/F ratio | Community Type | Associated herbs |
|---|---|---|---|---|---|---|---|---|
| ***M. aculeata*** | Rohtang Pass | 0.2 | 0.014 | 10.00 | 2.0 | 0.2 | *Rhododendron anthopogon* | *Polygonum paronychioides, Rhodiola quadrifida, Potentilla fulgens, Rhodiola heterodonta* |
| | Near Rani Nala, Rohtang Pass | 0.45 | 0.022 | 20.00 | 2.25 | 0.11 | *Anaphalis triplinervis-Poa alpina-Polygonatum verticillatum* | *Anaphalis triplinervis, Lagotis cashmeriana, Poa alpina, Polygonatum verticillatum* |
| ***R. campanulatum*** | Fozal, Beas valley | 110 | 0.21 | 75.00 | 1.46 | 0.01 | *Abies pindrow-Quercus semecarpifolia* | *Rosa macrophylla, Desmodium elegans, Rumex acetosa, Fragaria vesca* |
| | Fozal, Beas valley | 65 | 0.18 | 35 | 1.85 | 0.05 | *Abies pindrow* | *Rosa moschata, Berberis aristata, Juniperus communis, Rhododendron anthopogon* |

[1] Density is expressed as **individuals $m^{-2}$** for herbaceous communities (*Meconopsis aculeata*) and **individuals $ha^{-1}$** for shrub communities (*Rhododendron campanulatum*), reflecting standard phytosociological sampling methods for different vegetation strata.

| | | | | | | | | |
|---|---|---|---|---|---|---|---|---|
| | Fozal, Beas valley | 230 | 0.34 | 65 | 3.53 | 0.05 | *Quercus semecarpifolia* | *Juniperus communis, Salix denticulata, Rosa webbiana, Rhodiola heterodonta* |
| | Beas valley | 365 | 0.35 | 100 | 3.65 | 0.03 | *Rhododendron anthopogon* | *Euphrasia simplex, Corydalis govanniana, Bistorta vivipara, Anaphalis busua* |
| | Parvati valley | 115 | 0.24 | 60 | 1.91 | 0.03 | *Abies pindrow* | *Rosa webbiana, Juniperus indica, Bergenia stracheyi, Geranium wallichianum* |
| | Parvati valley | 110 | 0.36 | 85 | 3.23 | 0.03 | *Abies pindrow* | *Rosa webbiana, Jasminum officinale, Corydalis govaniana* |
| | Parvati valley | 150 | 0.32 | 55 | 2.72 | 0.04 | *Taxus contorta* | *Rosa webbiana, Cirsium wallichii, Jasminum officinale* |
| | Parvati valley | 185 | 0.35 | 65 | 2.84 | 0.04 | *Betula utilis* | *Rosa webbiana, Berberis glaucocarpa, Polygonatum verticillatum, Fragaria nubicola* |

| | | | | | | | | |
|---|---|---|---|---|---|---|---|---|
| | Parvati valley | 195 | 4.48 | 55 | 3.54 | 0.06 | *Betula utilis* | *Rhododendron lepidotum, Rosa webbiana, Lonicera asperifolia, Bergenia stracheyi* |
| | Jalori pass, Banjar | 3015 | 0.35 | 100 | 30.15 | 0.30 | *Rhododendron campanulatum* | *Rhododendron anthopogon, Rosa macrophylla, Cotoneaster affinis, Anaphalis adnata* |
| | Near Lambri top | 1235 | 0.31 | 100 | 12.35 | 0.12 | *Betula utilis* | *Rosa macrophylla, Cotoneaster microphyllus, Anaphalis busua, Parnassia nubicola* |
| | Lambri top, Sarchi | 1235 | 0.34 | 100 | 12.35 | 0.12 | *Rhododendron campanulatum* | *Ribes rubrum, Androsace rotundifolia, Achillea millefolium, Juncus concinnus* |
