## Supplementary files_2 for "Projected habitat loss and spatial redistribution of two alpine and subalpine species in Himachal Pradesh, Western Himalaya under CMIP6 scenarios": ESM_2.pdf

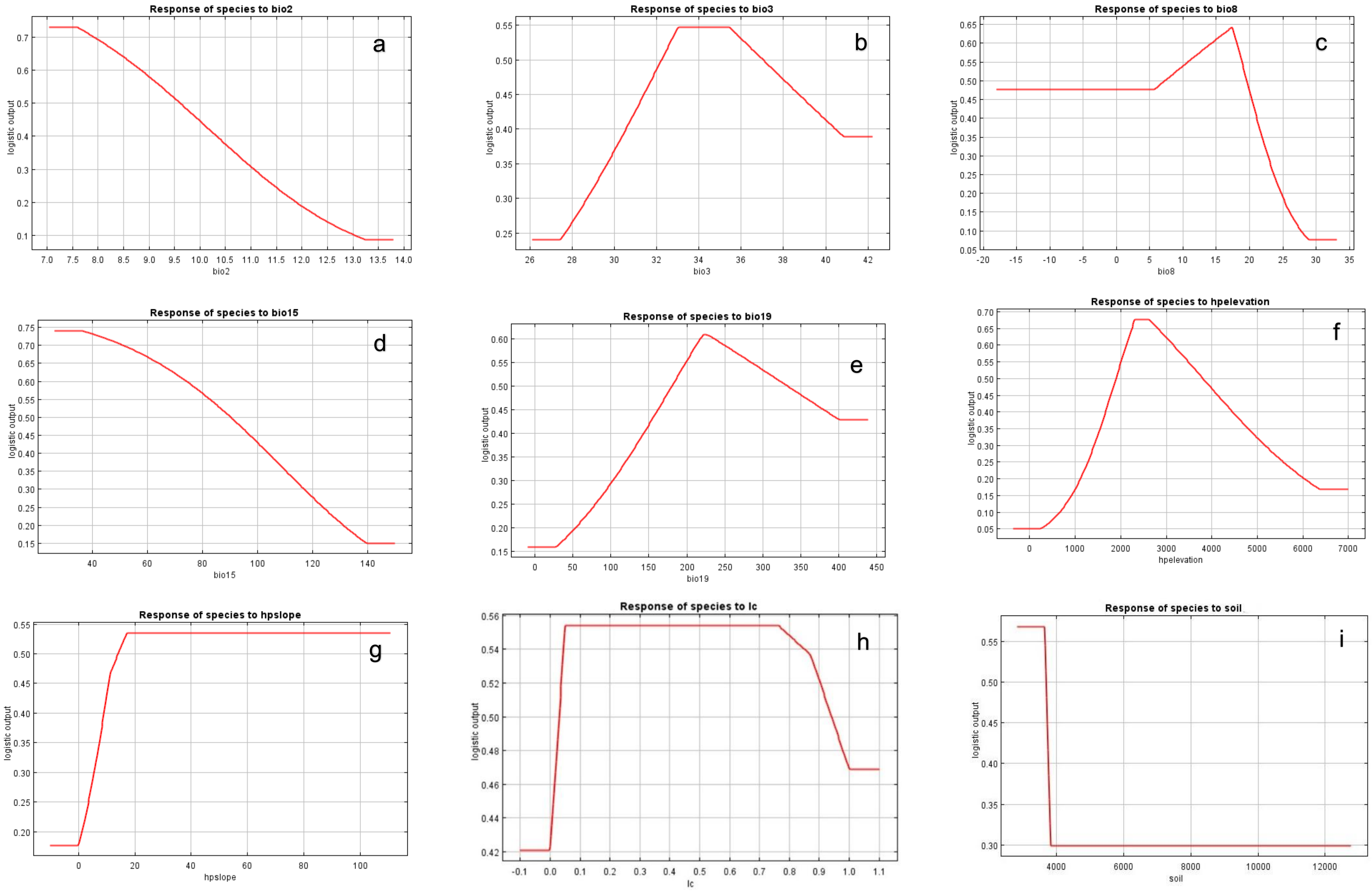
Response of species to bio2
a
logistic output
bio2
Response of species to bio3
b
logistic output
bio3
Response of species to bio8
c
logistic output
bio8
Response of species to bio15
d
logistic output
bio15
Response of species to bio19
e
logistic output
bio19
Response of species to hpelevation
f
logistic output
hpelevation
Response of species to hpslope
g
logistic output
hpslope
Response of species to lc
h
logistic output
lc
Response of species to soil
i
logistic output
soil

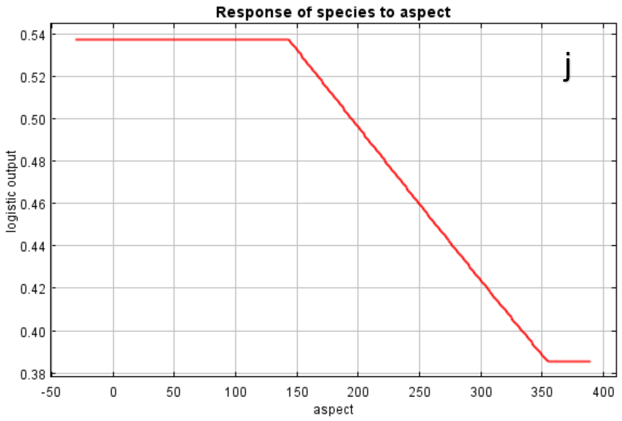


**Fig. S2** Response curves of MaxEnt-predicted habitat suitability (logistic output) for *M. aculeata* along the environmental gradients included in the final model. (a) Bio 2; Mean Diurnal Range (b) Bio 3; Isothermality, (c) Bio 8; Mean Temperature of Wettest Quarter, (d) Bio15; Precipitation seasonality, (e) Bio19; Precipitation of Coldest Quarter, (f) Elevation, (g) Slope, (h) Landcover, (i) Soil cover, (j) Aspect, in the species. The y-axis represents predicted habitat suitability (MaxEnt logistic output), and the x-axis represents the corresponding predictor value. For categorical land-cover data, x-axis values represent land-cover class codes and should not be interpreted as continuous measurements.

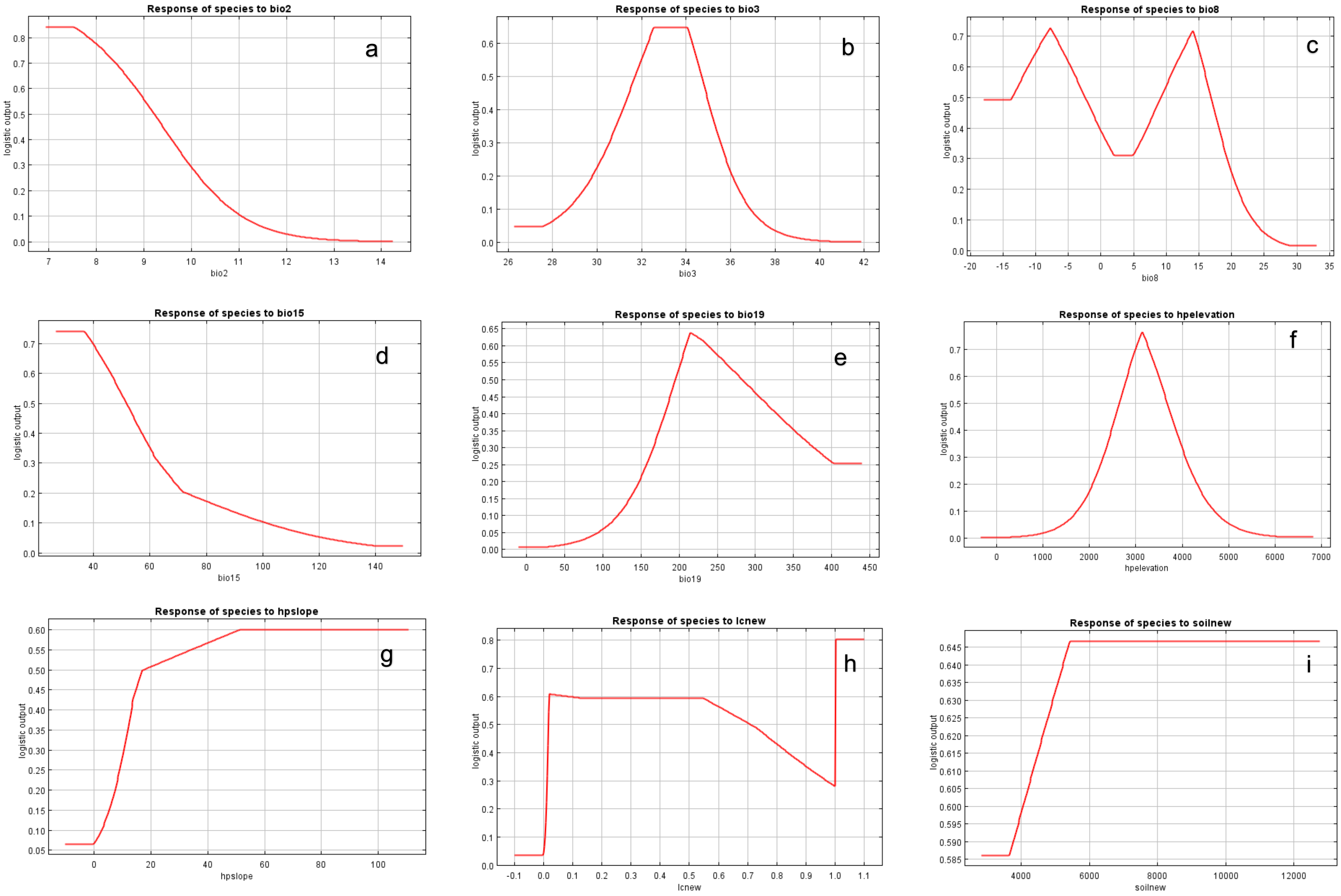
Response of species to bio2
a
logistic output
bio2
Response of species to bio3
b
logistic output
bio3
Response of species to bio8
c
logistic output
bio8
Response of species to bio15
d
logistic output
bio15
Response of species to bio19
e
logistic output
bio19
Response of species to hpelevation
f
logistic output
hpelevation
Response of species to hpslope
g
logistic output
hpslope
Response of species to lcnew
h
logistic output
lcnew
Response of species to soilnew
i
logistic output
soilnew

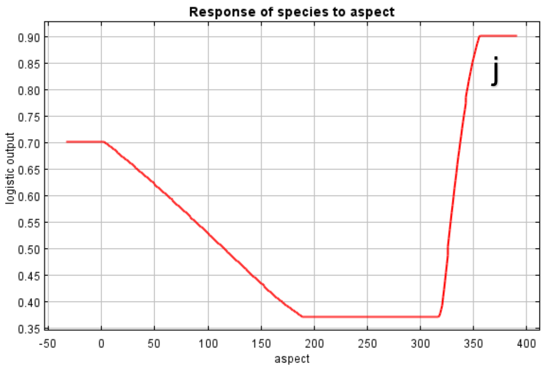


**Fig. S3** Response curves of MaxEnt-predicted habitat suitability (logistic output) for *R. campanulatum* along the environmental gradients included in the final model. (a) Bio 2; Mean Diurnal Range (b) Bio 3; Isothermality, (c) Bio 8; Mean Temperature of Wettest Quarter, (d) Bio15; Precipitation seasonality, (e) Bio19; Precipitation of Coldest Quarter, (f) Elevation, (g) Slope, (h) Landcover, (i) Soil cover, (j) Aspect, in the species. The y-axis represents predicted habitat suitability (MaxEnt logistic output), and the x-axis represents the corresponding predictor value. For categorical land-cover data, x-axis values represent land-cover class codes and should not be interpreted as continuous measurements.

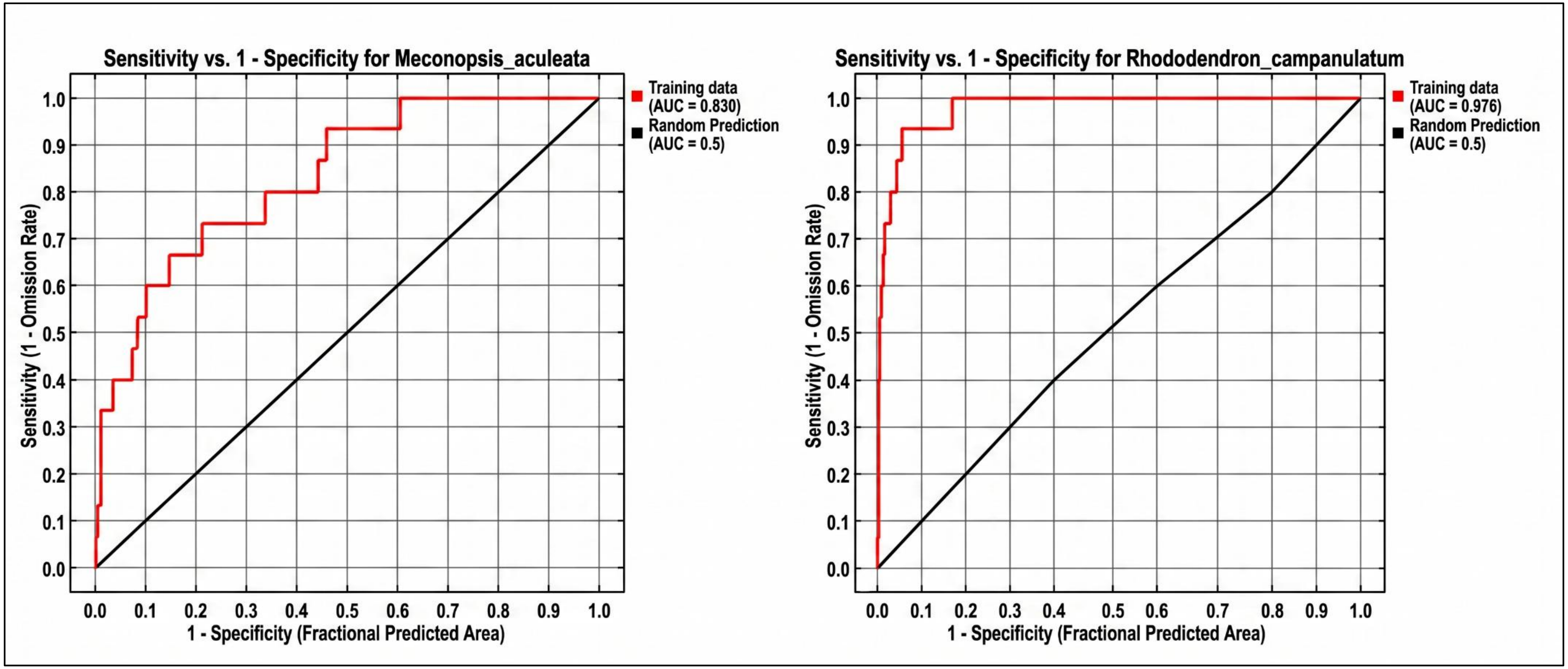


**Fig. S4** Receiver operating characteristic curve with area under curve (AUC) for *M. aculeata* (left panel) and *R. campanulatum* (right panel)

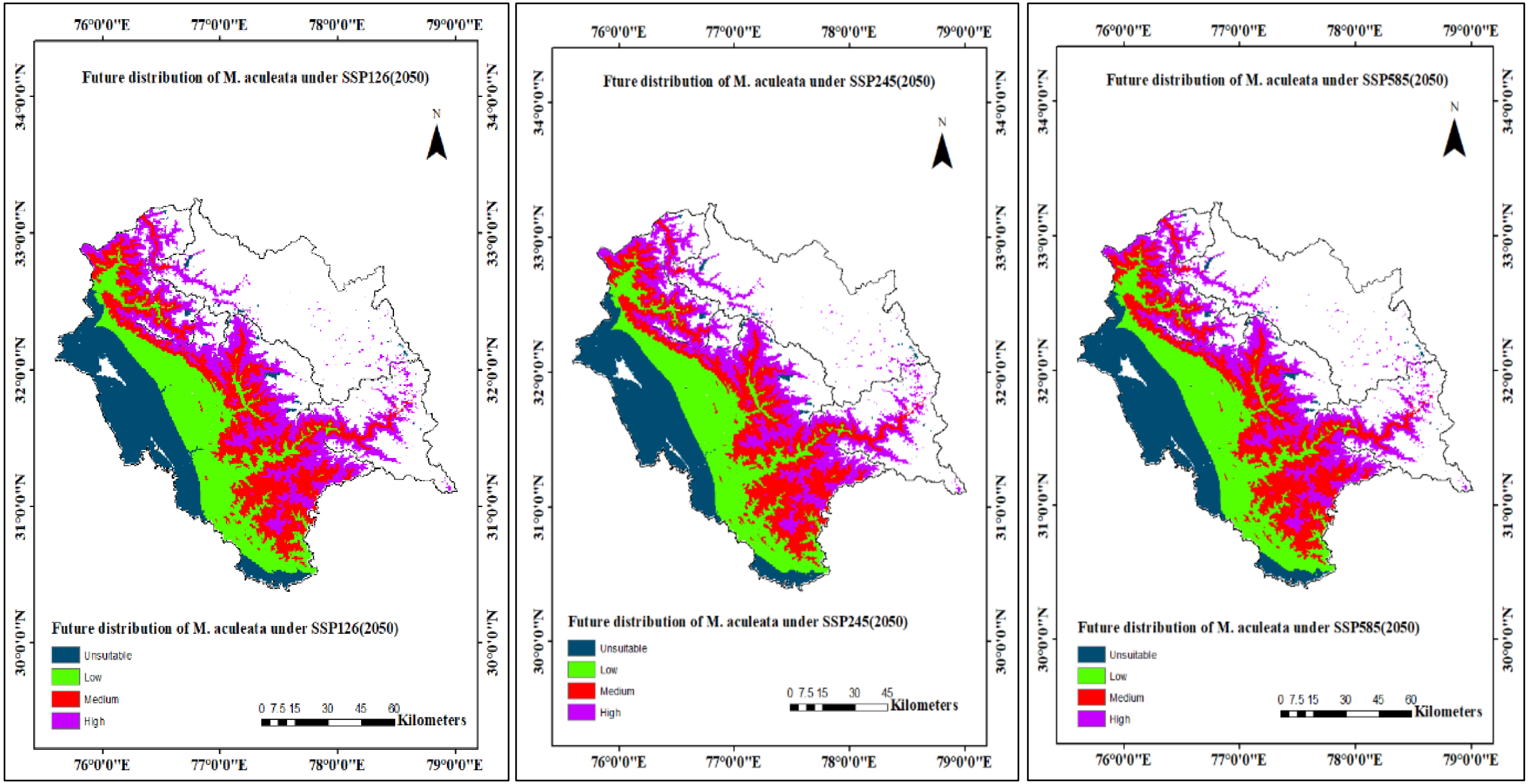
Future distribution of M. aculeata under SSP126(2050)
Future distribution of M. aculeata under SSP126(2050)
Unsuitable
Low
Medium
High
0 7.5 15 30 45 60
Kilometers
76°0'0"E
77°0'0"E
78°0'0"E
79°0'0"E
34°0'0"N
33°0'0"N
32°0'0"N
31°0'0"N
30°0'0"N
N
Fture distribution of M. aculeata under SSP245(2050)
Future distribution of M. aculeata under SSP245(2050)
Unsuitable
Low
Medium
High
0 7.5 15 30 45
Kilometers
Future distribution of M. aculeata under SSP585(2050)
Future distribution of M. aculeata under SSP585(2050)
Unsuitable
Low
Medium
High
0 7.5 15 30 45 60
Kilometers

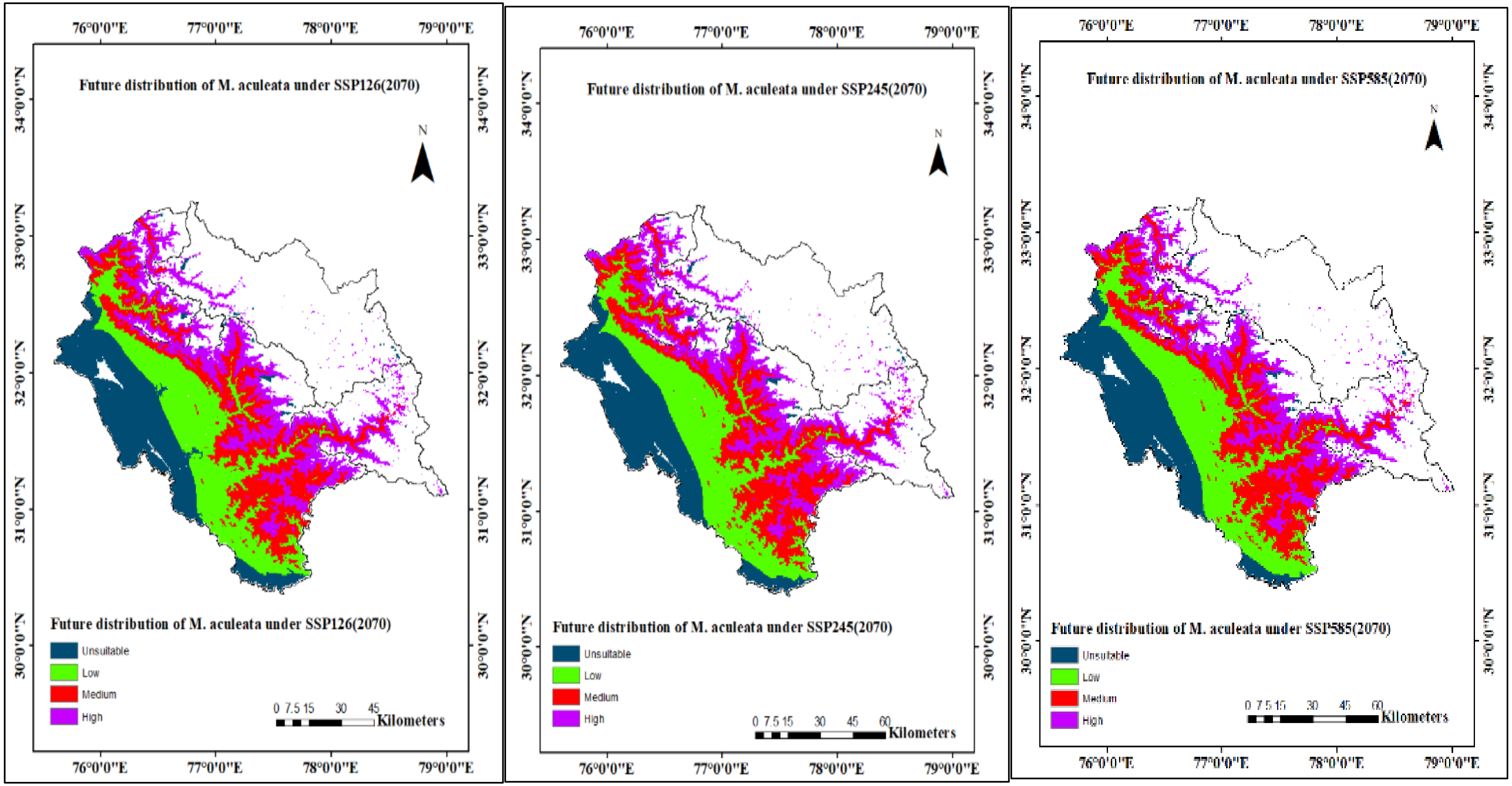


**Fig. S5** Future habitat suitability of *M. aculeata* based on future climate conditions; Blue region indicates no habitat suitability, green indicates poor habitat suitability, red indicates fair habitat suitability, purple indicates good habitat suitability. Maxent 3.4.3 (https://biodiversityinformatics.amnh.org/open_source/maxent/); ArcMap 10.8 (https://unikassel-its.maps.arcgis.com/home).

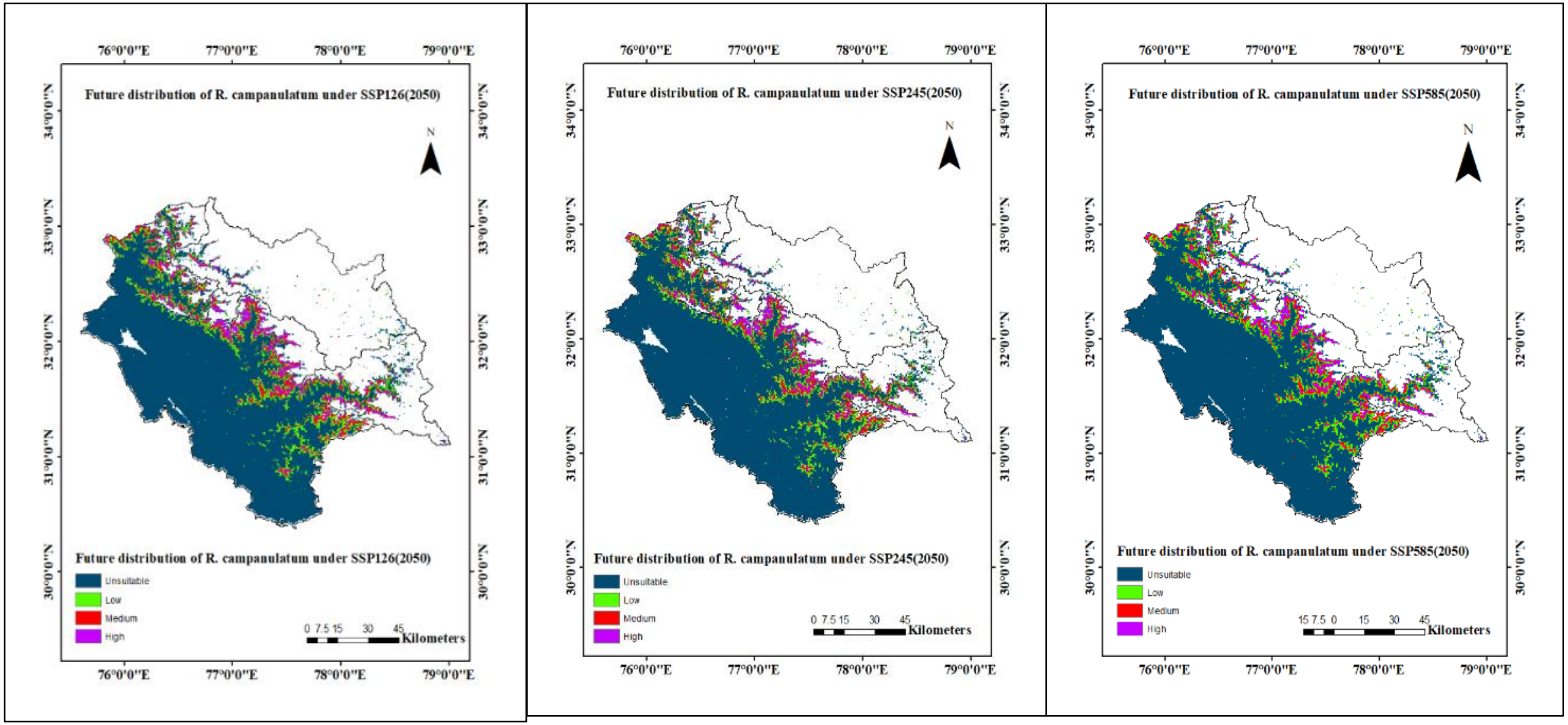
Future distribution of R. campanulatum under SSP126(2050)
Future distribution of R. campanulatum under SSP126(2050)
Unsuitable
Low
Medium
High
0 7.5 15 30 45
Kilometers
76°0'0"E
77°0'0"E
78°0'0"E
79°0'0"E
34°0'0"N
33°0'0"N
32°0'0"N
31°0'0"N
30°0'0"N
N
Future distribution of R. campanulatum under SSP245(2050)
Future distribution of R. campanulatum under SSP245(2050)
Unsuitable
Low
Medium
High
0 7.5 15 30 45
Kilometers
Future distribution of R. campanulatum under SSP585(2050)
Future distribution of R. campanulatum under SSP585(2050)
Unsuitable
Low
Medium
High
15 7.5 0 15 30 45
Kilometers

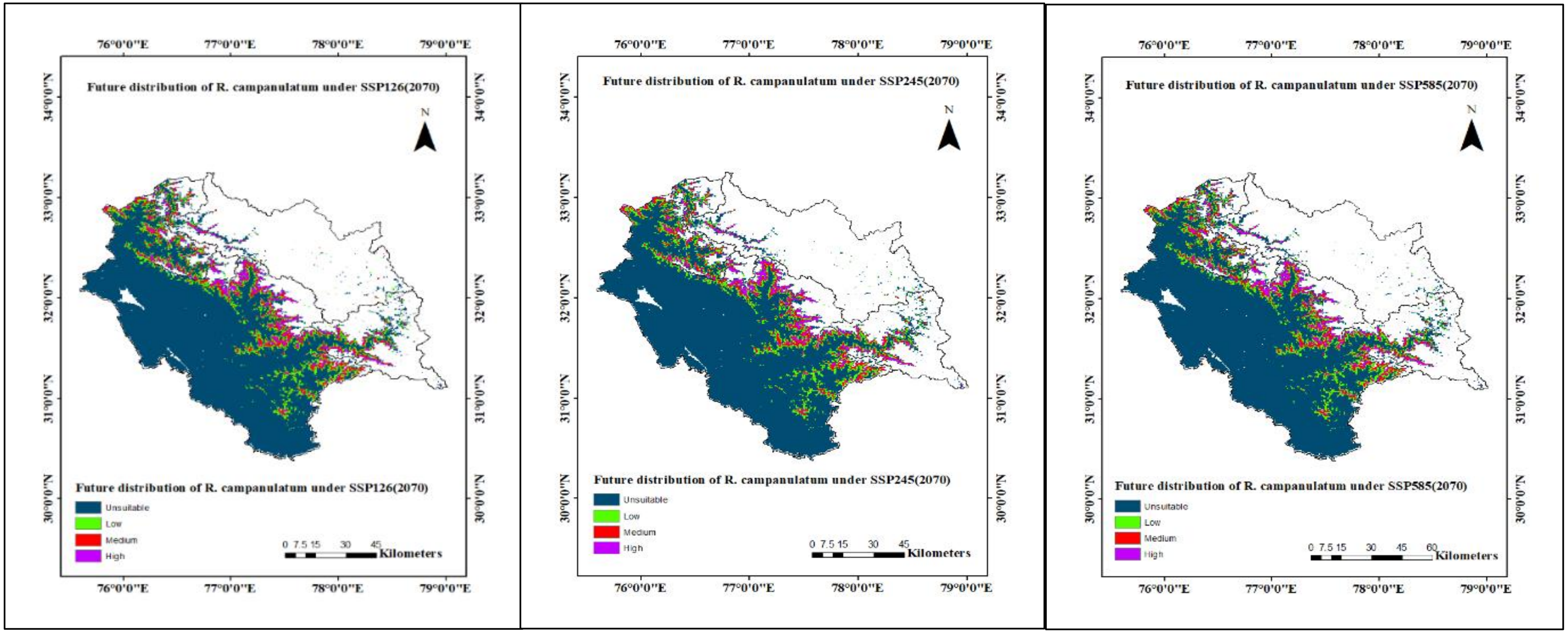


**Fig. S6** Future habitat suitability of *R. campanulatum* based on future climate conditions; Blue region indicates no habitat suitability, green indicates poor habitat suitability, red indicates fair habitat suitability, purple indicates good habitat suitability. Maxent 3.4.3 (https://biodiversityinformatics.amnh.org/open_source/maxent/); ArcMap 10.8 (https://unikassel-its.maps.arcgis.com/home).

**Table S7** Predicted suitable areas ($km^2$) under both current and future climatic conditions

| Species | Scenario | Year | Area (in km2) | | | | | | |
|---|---|---|---|---|---|---|---|---|---|
| | | | Unsuitable | Low (25-50%) | Medium (50-75%) | High (>75%) | Total suitable area | Loss (-) or Gain (+) | (%) Increase/ decrease |
| *M. aculeata* | Current | | 1649.51 | 1160.32 | 744.33 | 881.17 | 2785.82 | - | - |
| | SSP126 | 2050 | 586.10 | 713.97 | 697.39 | 665.07 | 2076.43 | (-) 709.39 | 25.46 |
| | | 2070 | 594.71 | 717.41 | 679.33 | 671.13 | 2067.87 | (-) 717.95 | 25.77 |
| | SSP245 | 2050 | 581.88 | 710.58 | 712.66 | 657.52 | 2080.76 | (-) 705.06 | 25.30 |
| | | 2070 | 579.26 | 717.77 | 729.18 | 636.31 | 2083.26 | (-) 702.56 | 25.21 |
| | SSP585 | 2050 | 579.56 | 713.73 | 728.23 | 641.12 | 2083.08 | (-) 702.74 | 25.22 |

| | | | | | | | | |
|---|---|---|---|---|---|---|---|---|
| | | 2070 | 572.61 | 713.43 | 765.90 | 610.76 | 2090.09 | (-)<br>695.73 | 24.97 |
| *R. campanulatum* | Current | | 3657.44 | 404.46 | 218.12 | 155.31 | 777.89 | - | - |
| | SSP126 | 2050 | 2101.92 | 271.18 | 166.37 | 123.05 | 560.60 | (-)<br>217.29 | 27.93 |
| | | 2070 | 2097.82 | 278.13 | 166.01 | 120.61 | 564.75 | (-)<br>213.14 | 27.39 |
| | SSP245 | 2050 | 2098.00 | 269.10 | 168.74 | 126.79 | 564.63 | (-)<br>213.26 | 27.14 |
| | | 2070 | 2104.48 | 270.23 | 165.24 | 122.58 | 558.05 | (-)<br>219.84 | 28.26 |
| | SSP585 | 2050 | 2099.19 | 270.65 | 167.73 | 125.07 | 563.45 | (-)<br>214.44 | 27.56 |
| | | 2070 | 2104.83 | 270.05 | 164.82 | 122.99 | 557.86 | (-)<br>220.03 | 28.28 |

**Table S8** Projected centroid displacement, movement direction (bearing), and climatic niche overlap (Schoener's D) for *M. aculeata* and *R. campanulatum* under future climate scenarios

| **Species** | **SSP** | **Year** | **Centroid shift (km)** | **Bearing (°)** | **Schoener's *D*** |
|---|---|---|---|---|---|
| *M. aculeata* | SSP126 | 2050 | 8.50 | 209.12 | 0.673417 |
| | | 2070 | 8.50 | 209.12 | 0.673492 |
| | SSP245 | 2050 | 8.50 | 209.12 | 0.673468 |
| | | 2070 | 8.50 | 209.12 | 0.673545 |
| | SSP585 | 2050 | 7.74 | 180.39 | 0.506886 |
| | | 2070 | 7.74 | 180.39 | 0.506886 |
| *R. campanulatum* | SSP126 | 2050 | 8.41 | 209.84 | 0.688908 |
| | | 2070 | 8.50 | 209.12 | 0.687905 |
| | SSP245 | 2050 | 8.50 | 209.12 | 0.687845 |
| | | 2070 | 8.50 | 209.12 | 0.687401 |
| | SSP585 | 2050 | 8.50 | 209.12 | 0.688051 |
| | | 2070 | 8.50 | 209.12 | 0.687761 |
