## Supplementary files_3 for "Projected habitat loss and spatial redistribution of two alpine and subalpine species in Himachal Pradesh, Western Himalaya under CMIP6 scenarios": ESM_3.pdf

**SSP 1-2.6 (2050)**

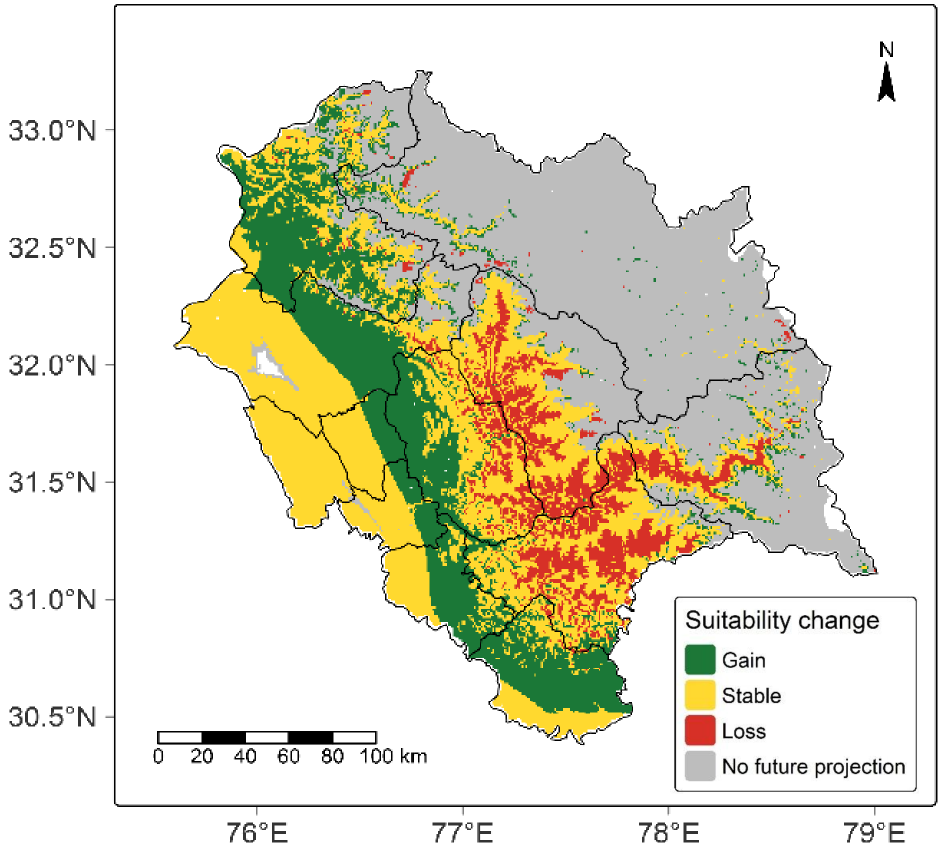


**SSP1-2.6 (2070)**

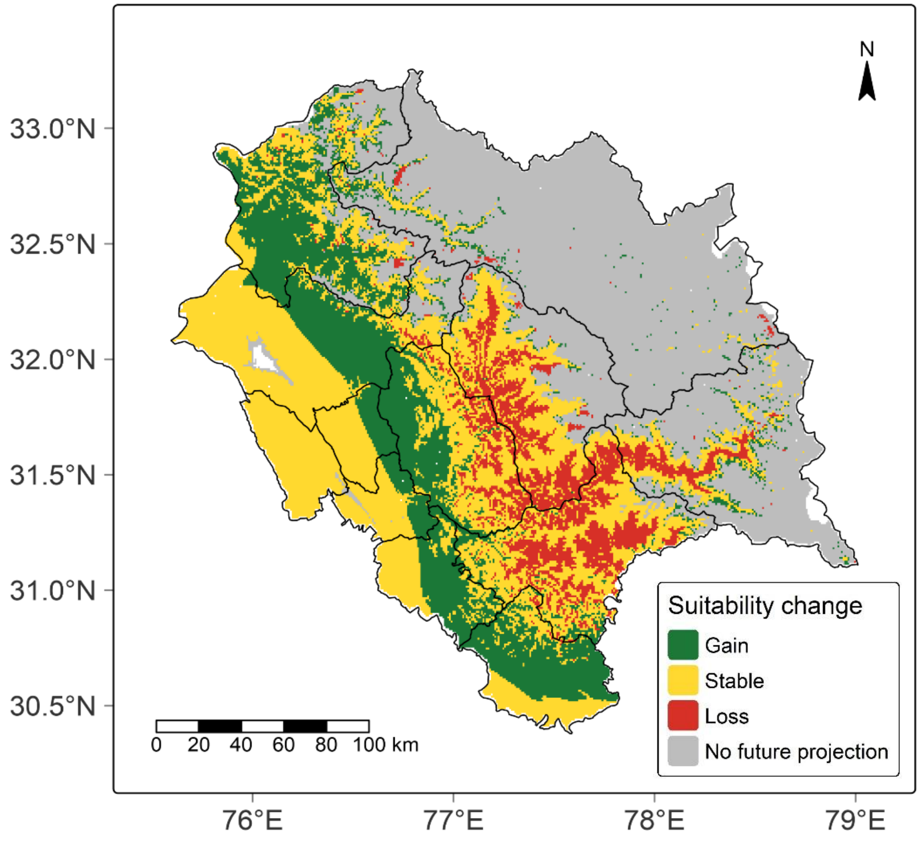

**SSP 2-4.5 (2050)**

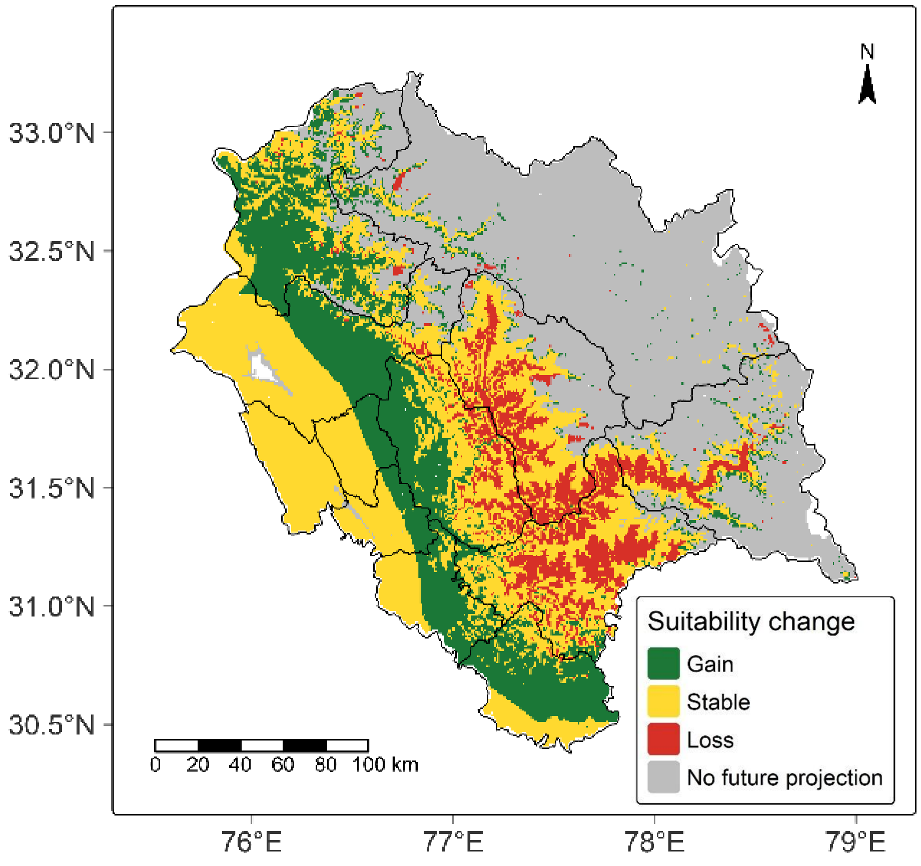


**SSP 2-4.5 (2070)**

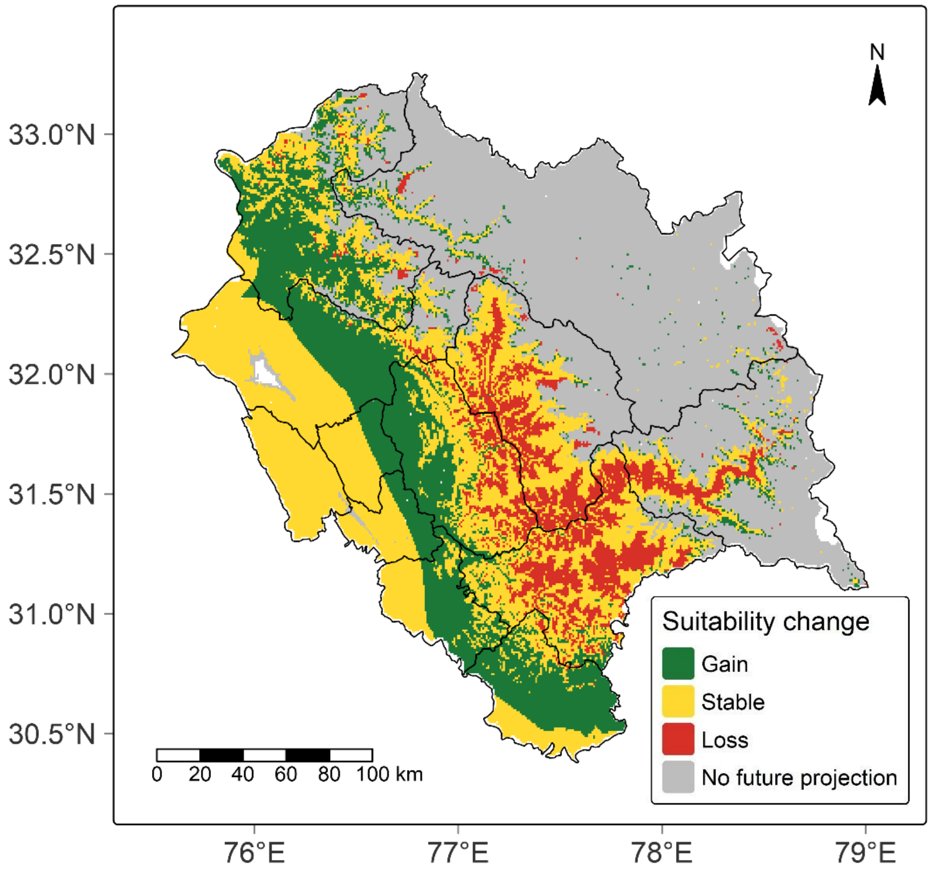

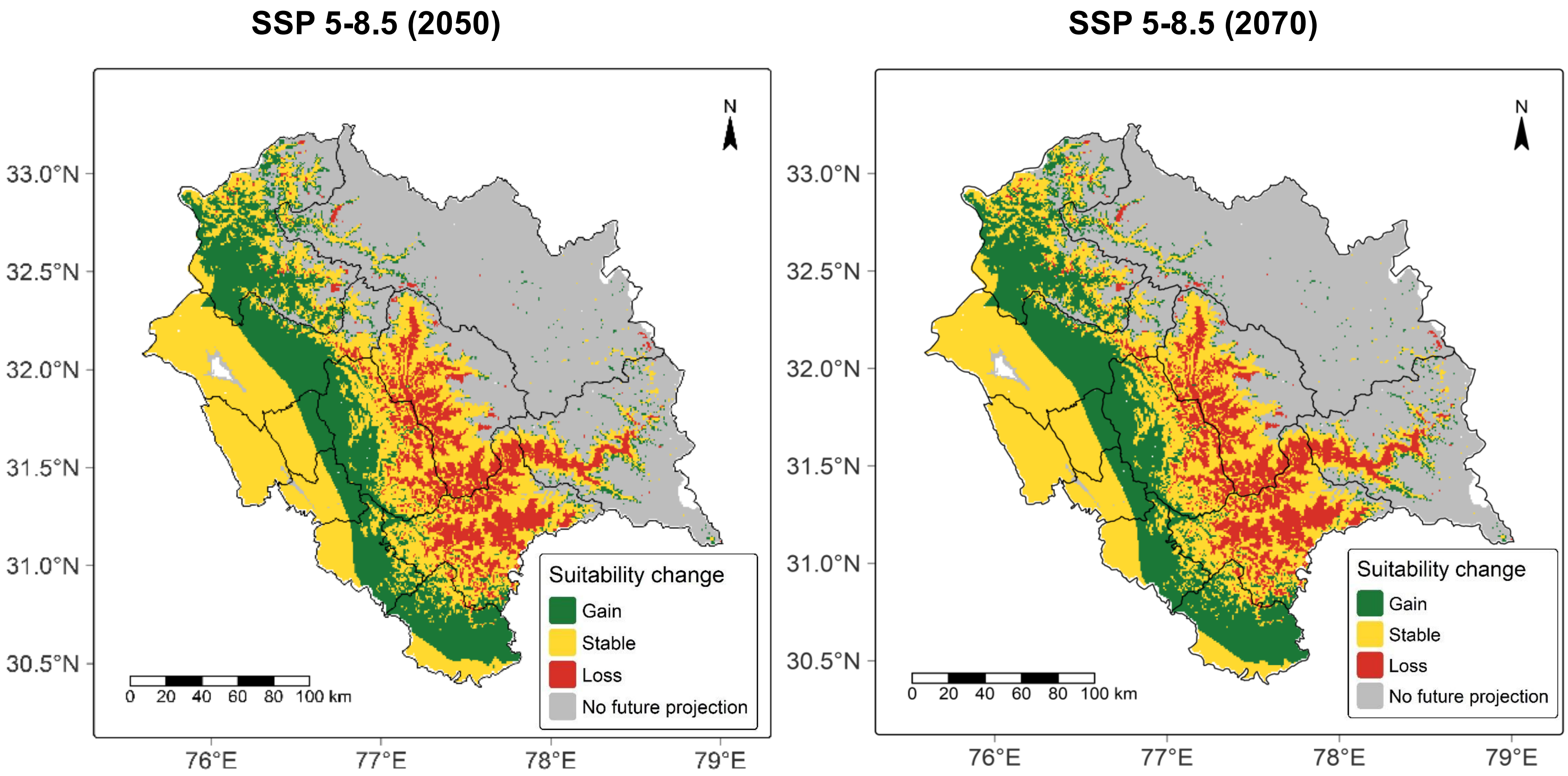


**Fig. S7** Suitability change maps for M. aculeata for SSP-1.2.6 and 2-4.5 for year 2050 and 2070

**SSP 1-2.6 (2050)**

**SSP 1-2.6 (2070)**

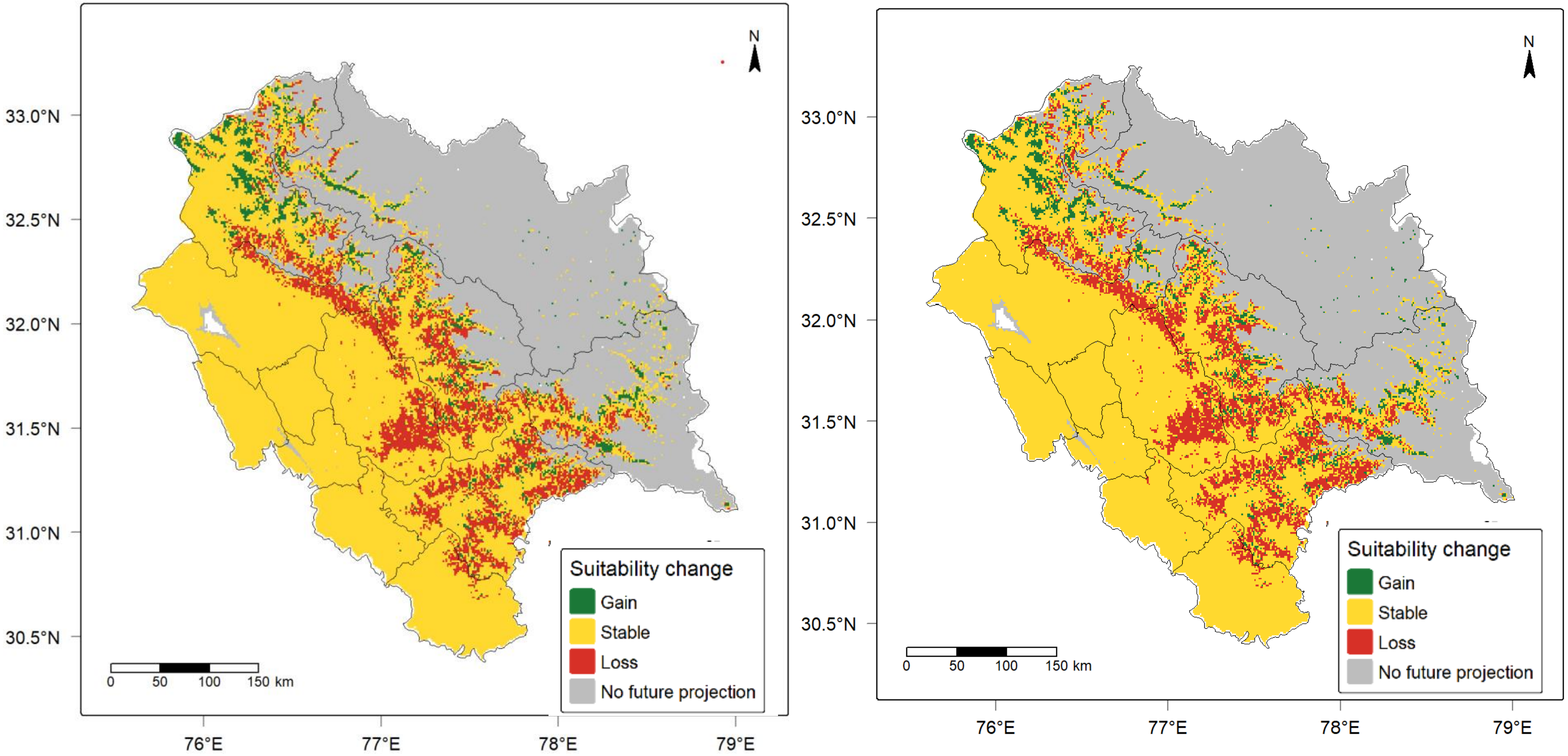

### SSP 2-4.5 (2050)

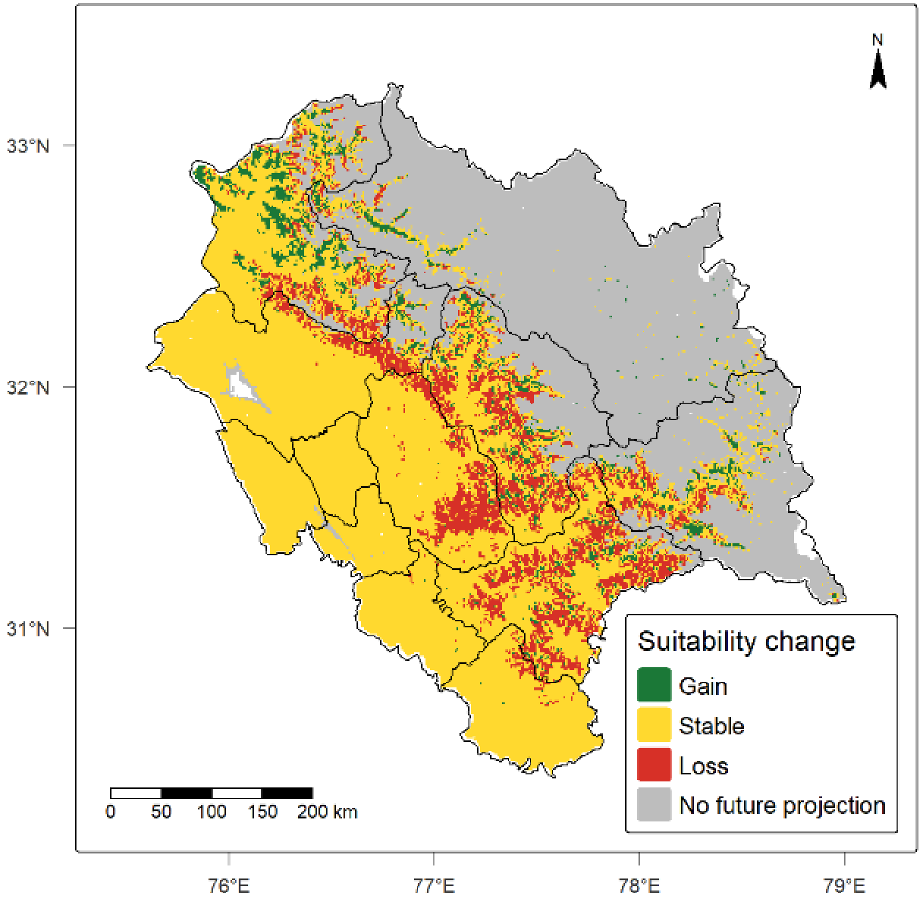


### SSP 2-4.5 (2070)

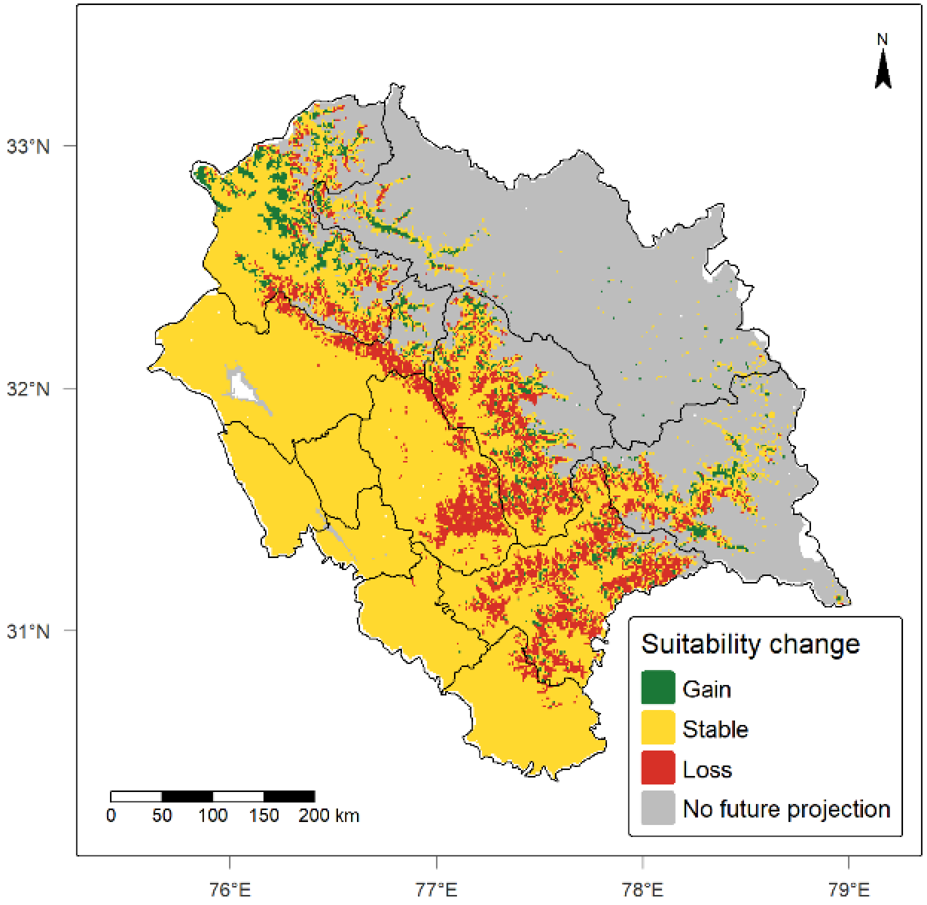

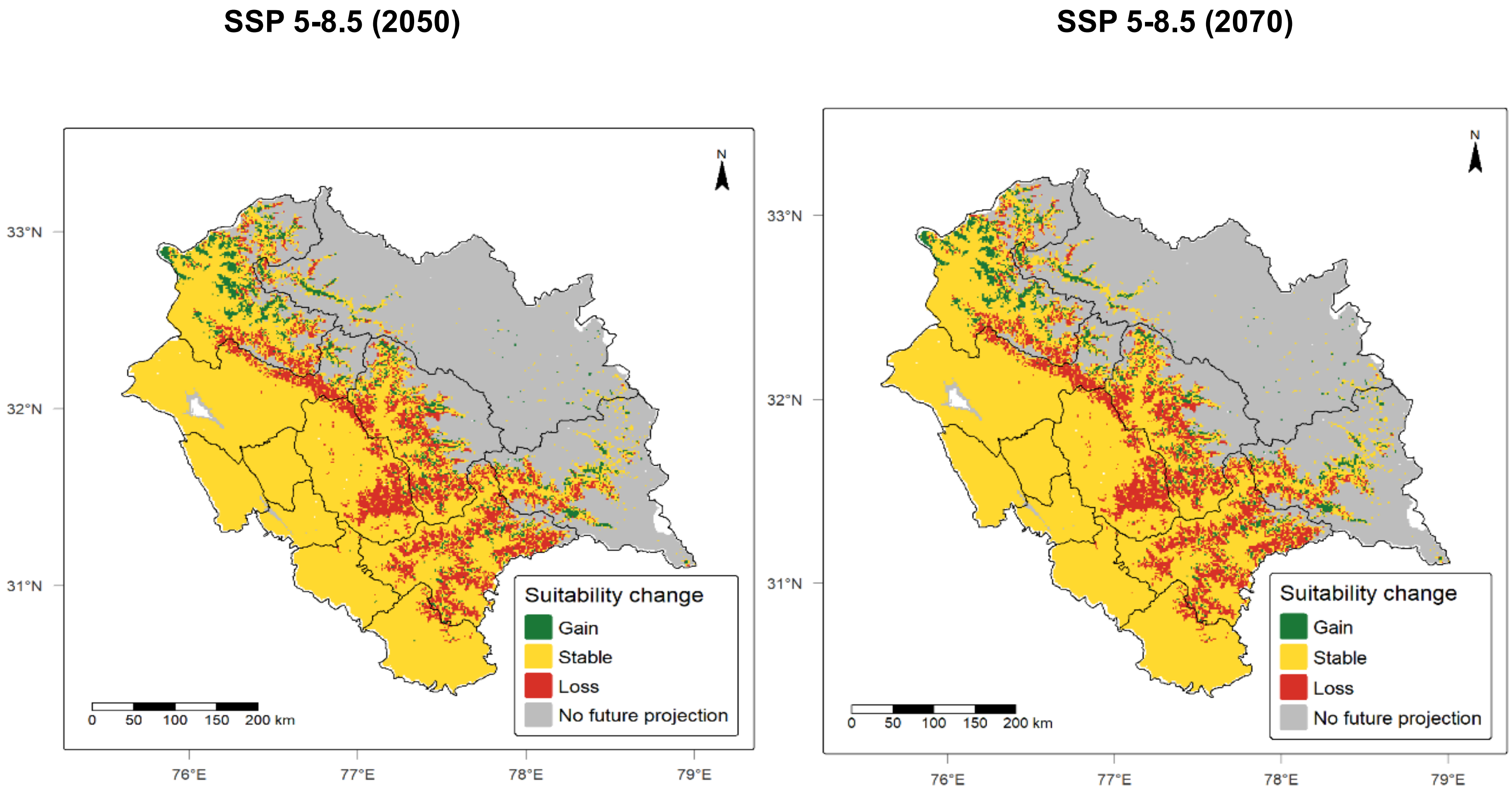


**Fig. S8** Suitability change maps for *R. campanulatum*

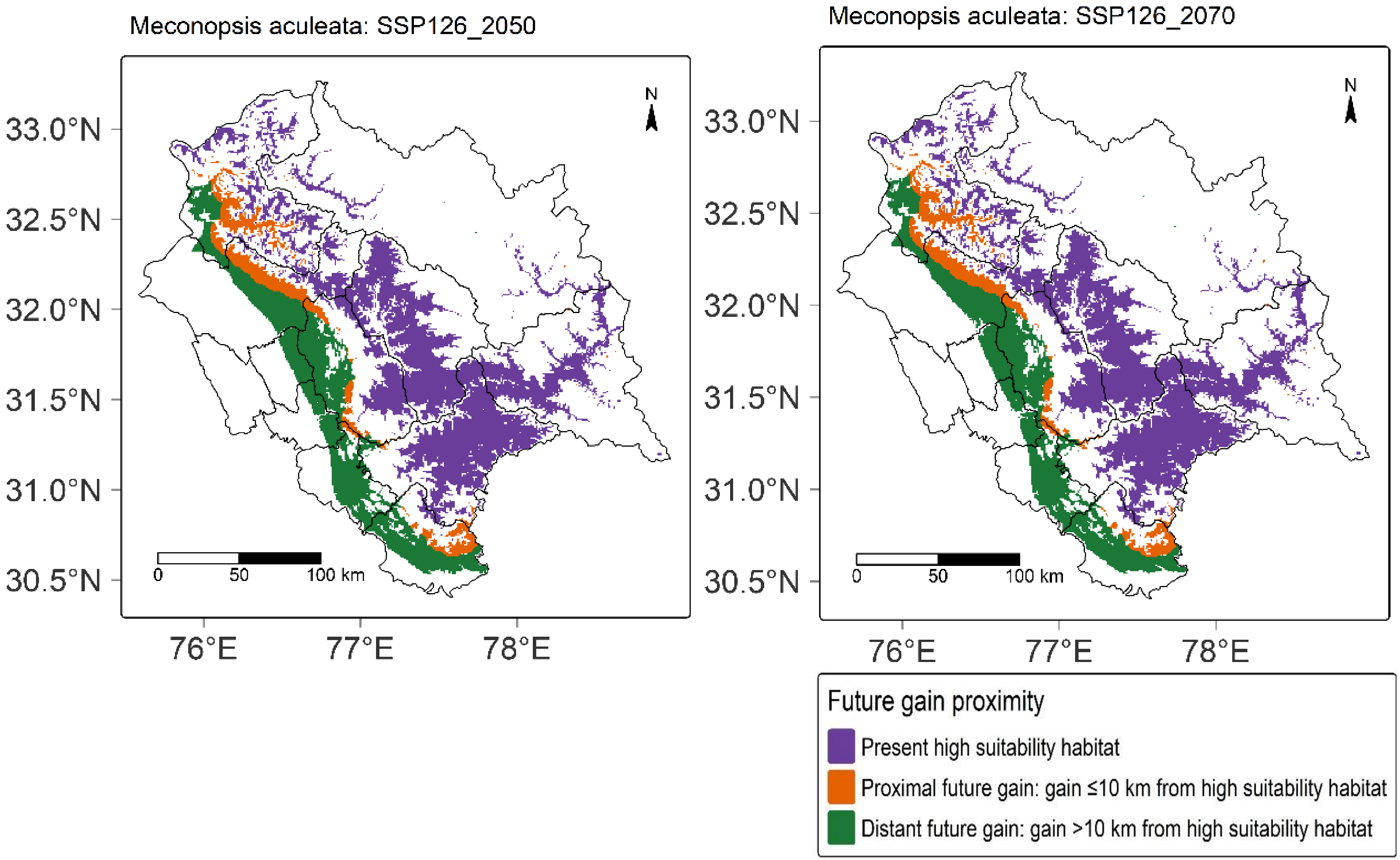
Meconopsis aculeata: SSP126_2050
Meconopsis aculeata: SSP126_2070
33.0°N
32.5°N
32.0°N
31.5°N
31.0°N
30.5°N
76°E
77°E
78°E
N
0
50
100 km
Future gain proximity
Present high suitability habitat
Proximal future gain: gain ≤10 km from high suitability habitat
Distant future gain: gain >10 km from high suitability habitat

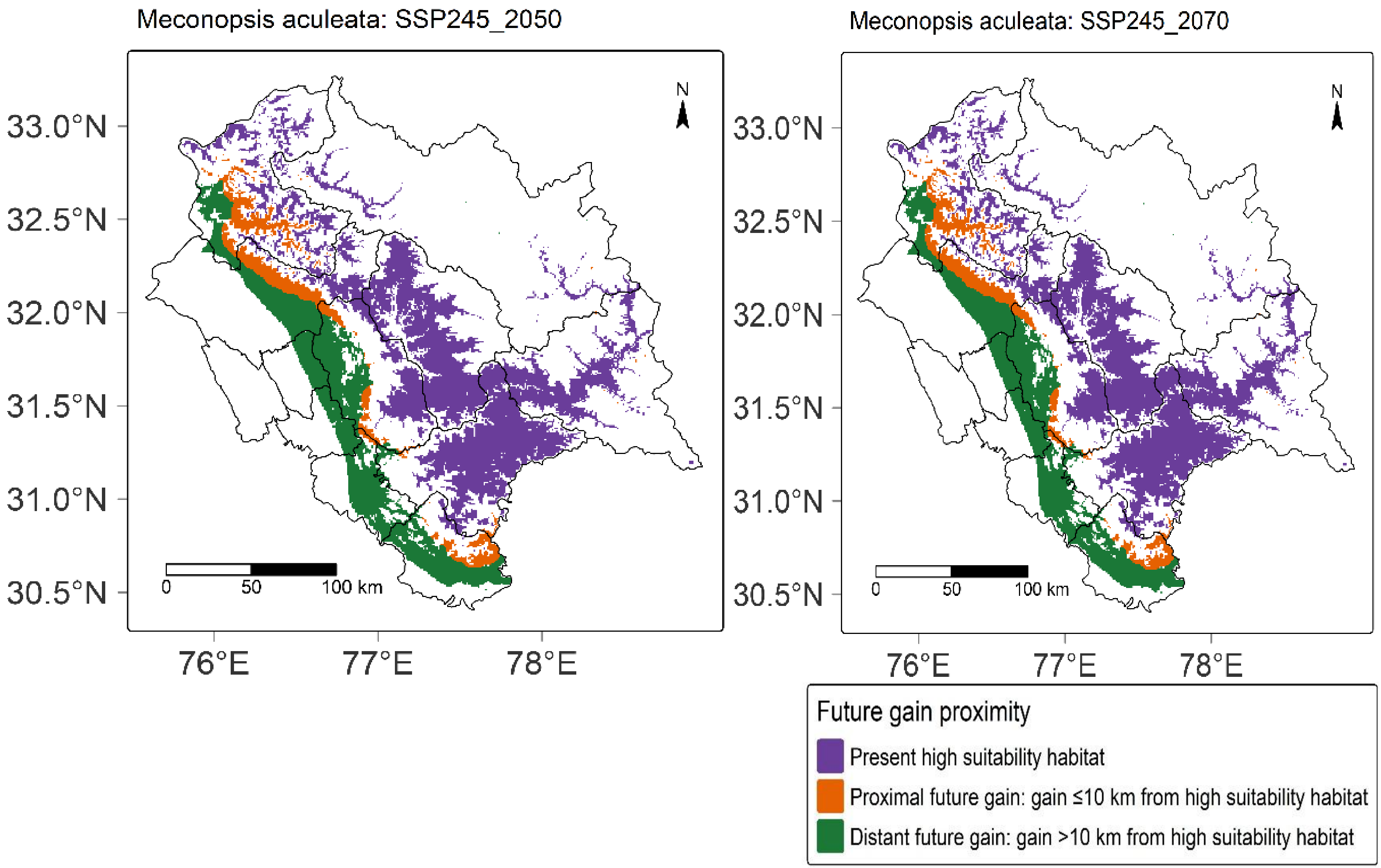
Meconopsis aculeata: SSP245_2050
Meconopsis aculeata: SSP245_2070
33.0°N
32.5°N
32.0°N
31.5°N
31.0°N
30.5°N
76°E
77°E
78°E
N
0
50
100 km
Future gain proximity
Present high suitability habitat
Proximal future gain: gain ≤10 km from high suitability habitat
Distant future gain: gain >10 km from high suitability habitat

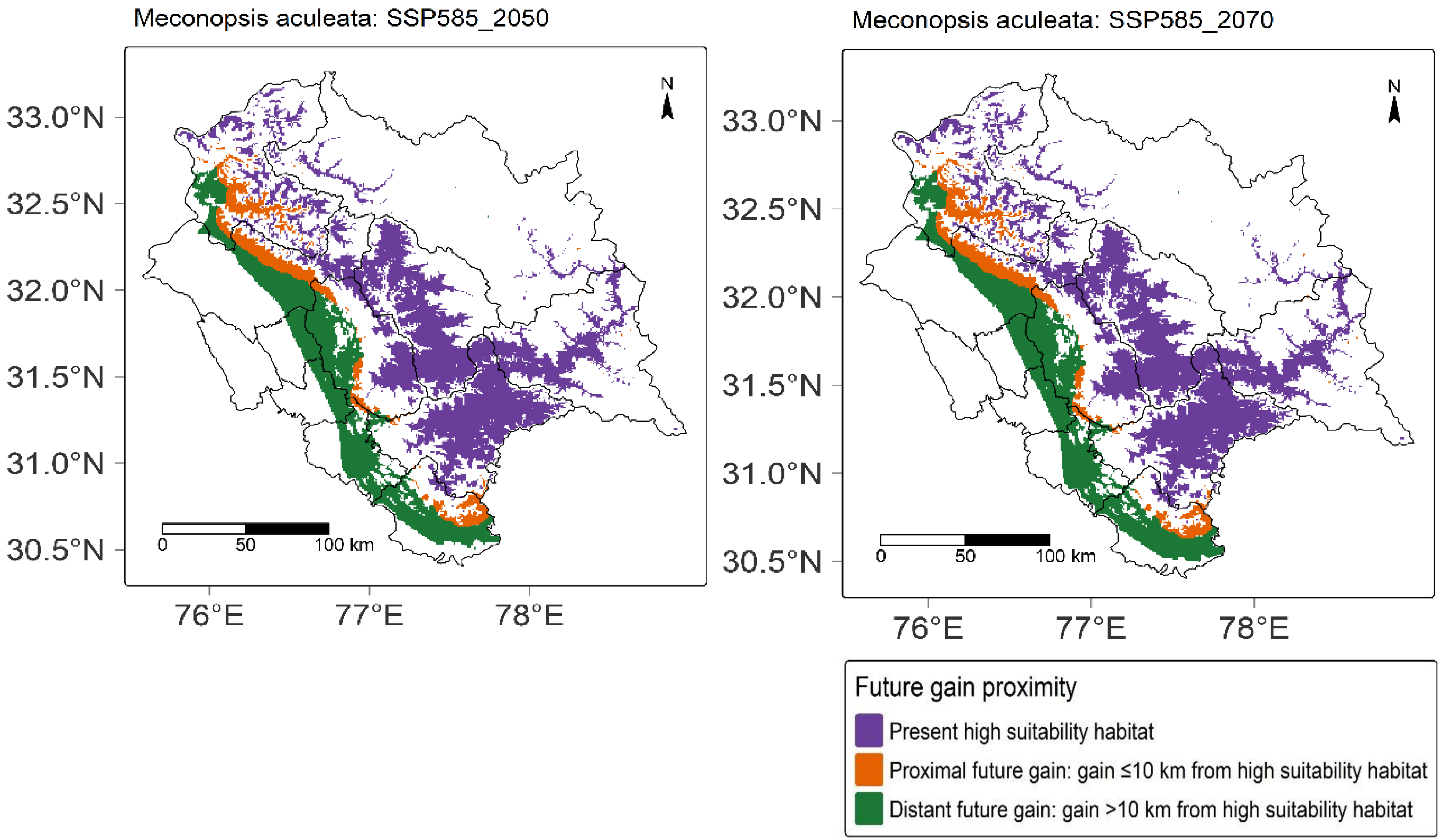


**Fig. S9** Proximity maps for *M. aculeata*

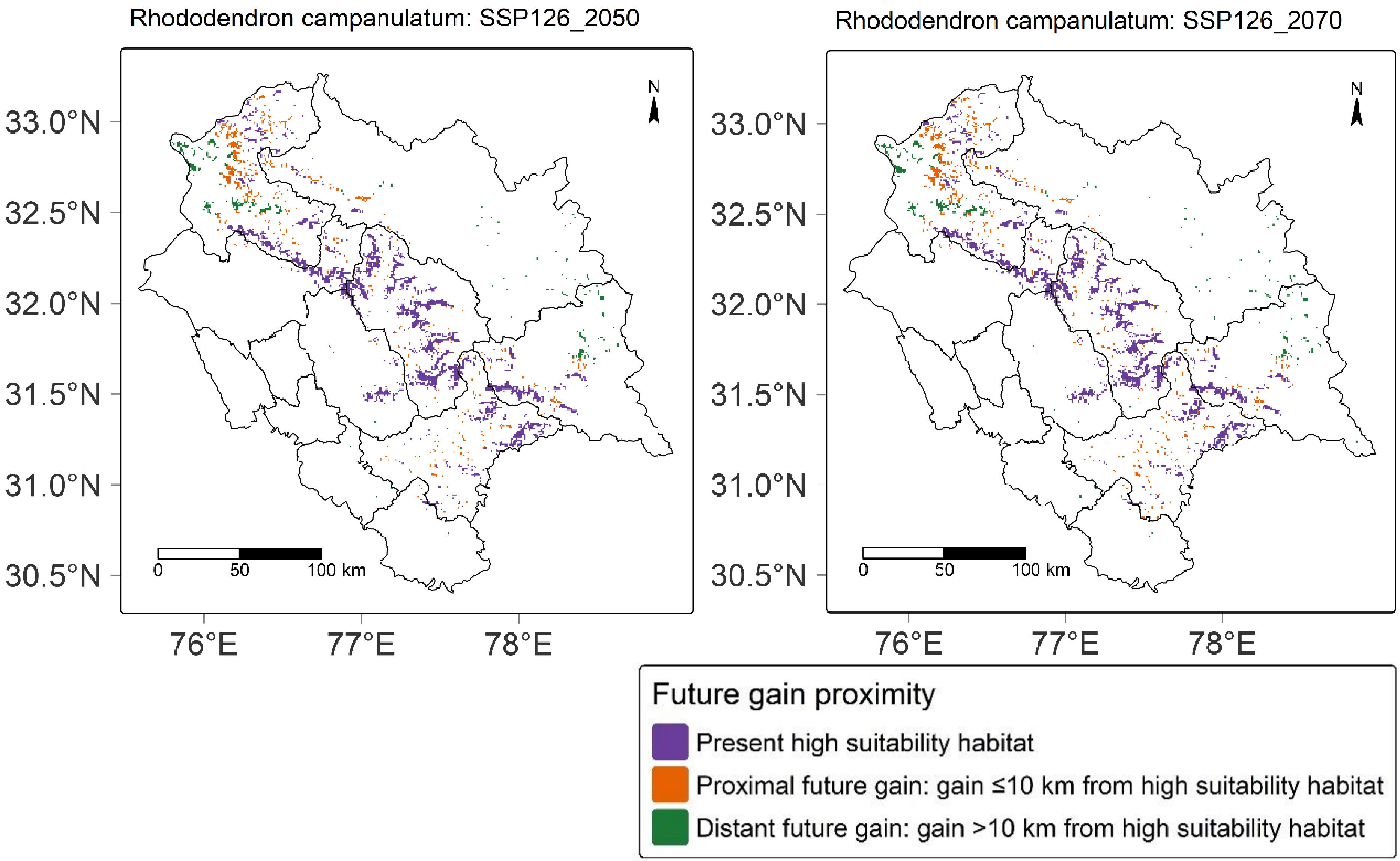

Rhododendron campanulatum: SSP126_2050
Rhododendron campanulatum: SSP126_2070
33.0°N
32.5°N
32.0°N
31.5°N
31.0°N
30.5°N
76°E
77°E
78°E
N
0
50
100 km
Future gain proximity
Present high suitability habitat
Proximal future gain: gain ≤10 km from high suitability habitat
Distant future gain: gain >10 km from high suitability habitat

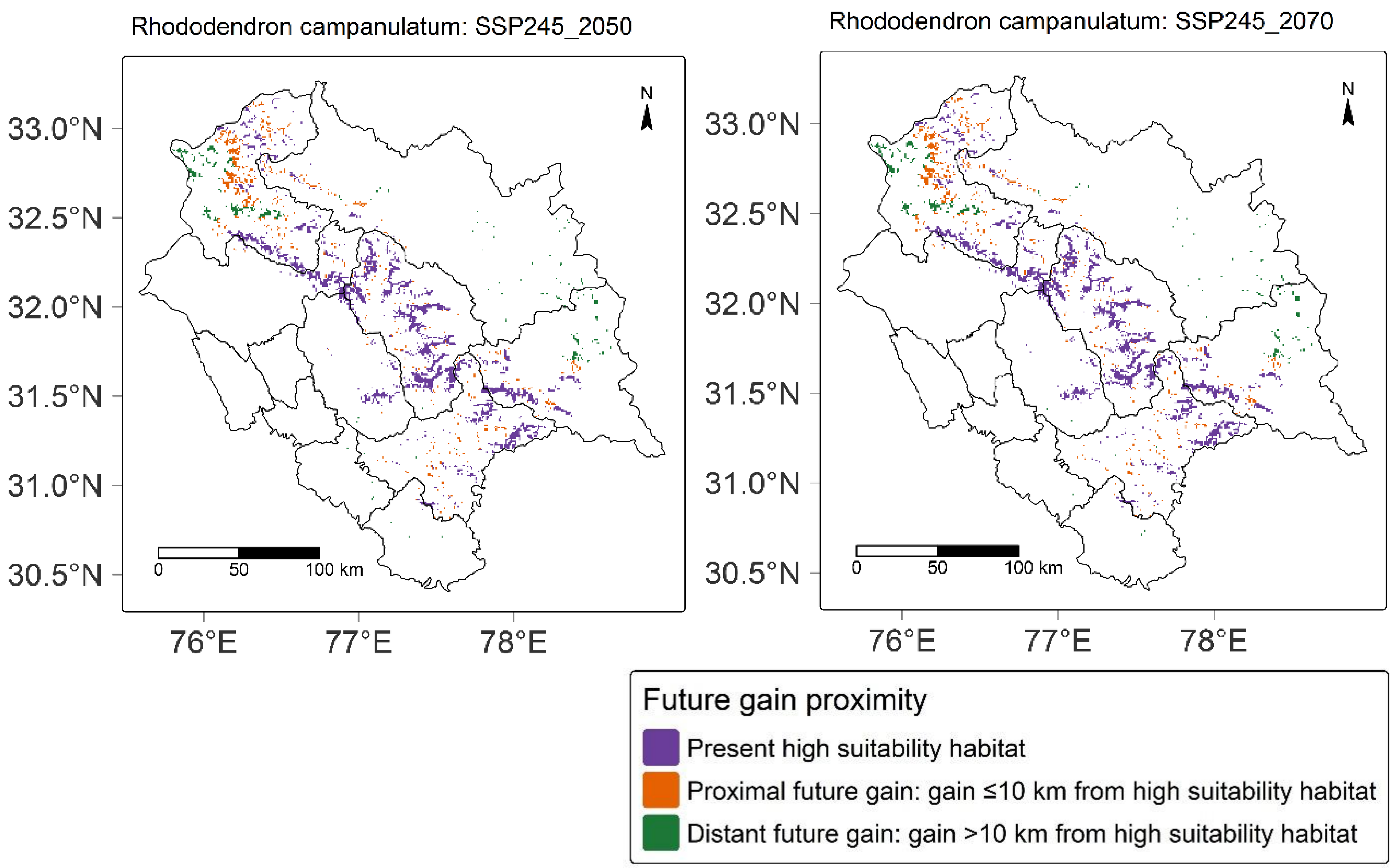
Rhododendron campanulatum: SSP245_2050
Rhododendron campanulatum: SSP245_2070
N
33.0°N
32.5°N
32.0°N
31.5°N
31.0°N
30.5°N
76°E
77°E
78°E
0
50
100 km
Future gain proximity
Present high suitability habitat
Proximal future gain: gain ≤10 km from high suitability habitat
Distant future gain: gain >10 km from high suitability habitat

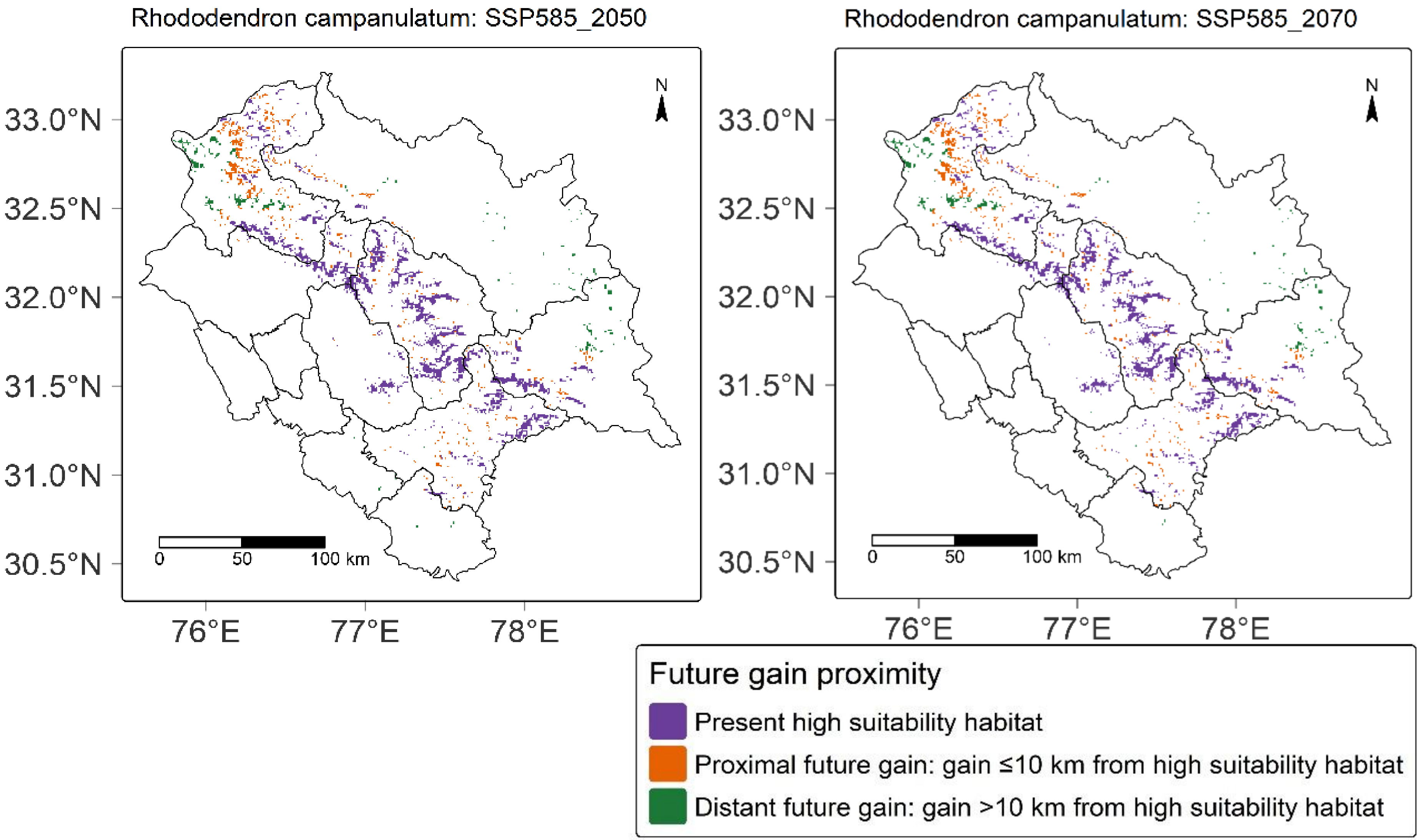


**Fig. S10** Proximity maps for *R. campanulatum*
