## Supplementary files_3 for "Projected habitat loss and spatial redistribution of two alpine and subalpine species in Himachal Pradesh, Western Himalaya under CMIP6 scenarios": ESM_4.pdf

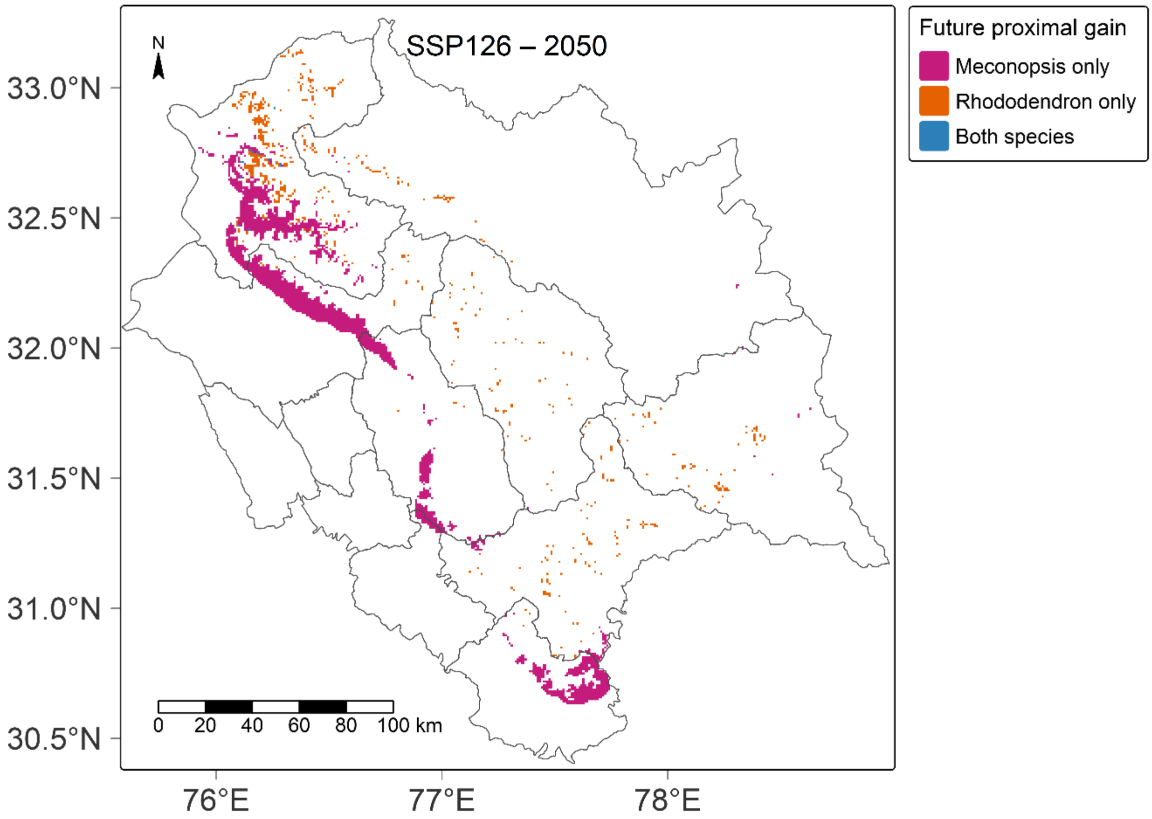

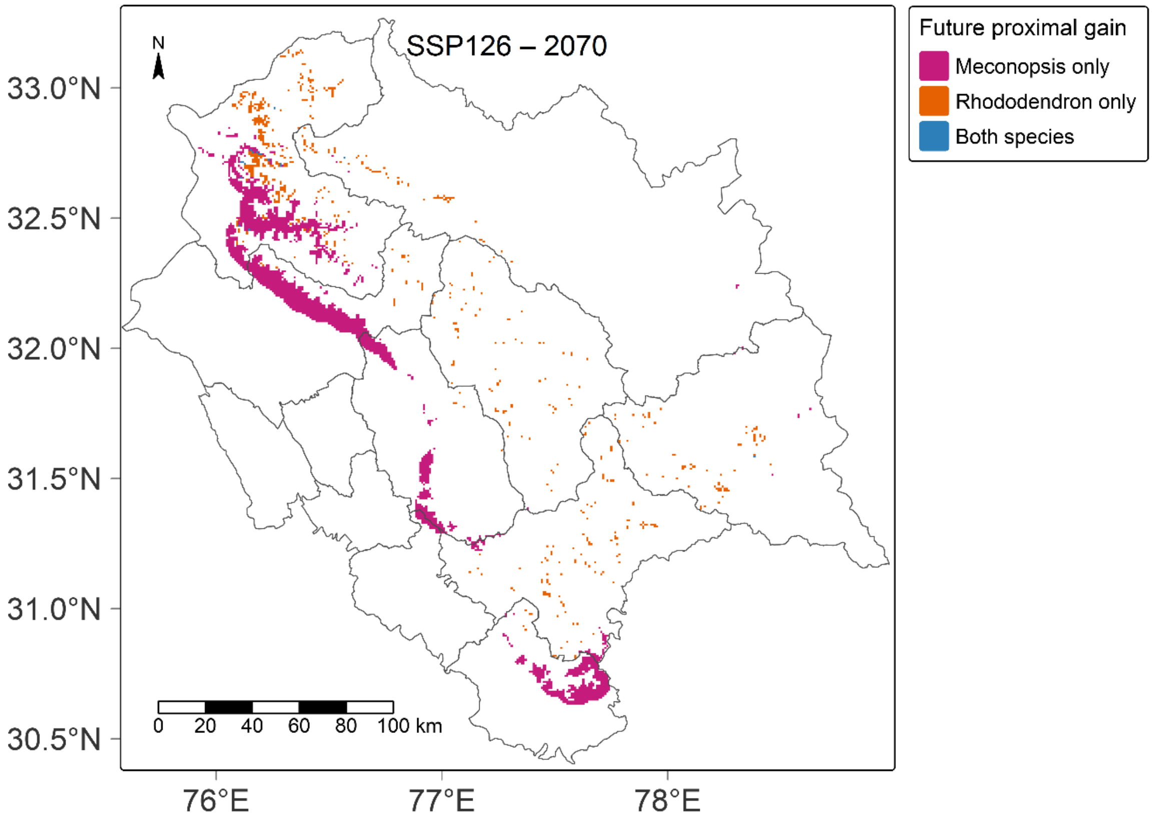
SSP126 – 2070
Future proximal gain
Meconopsis only
Rhododendron only
Both species
N
0 20 40 60 80 100 km
33.0°N
32.5°N
32.0°N
31.5°N
31.0°N
30.5°N
76°E
77°E
78°E

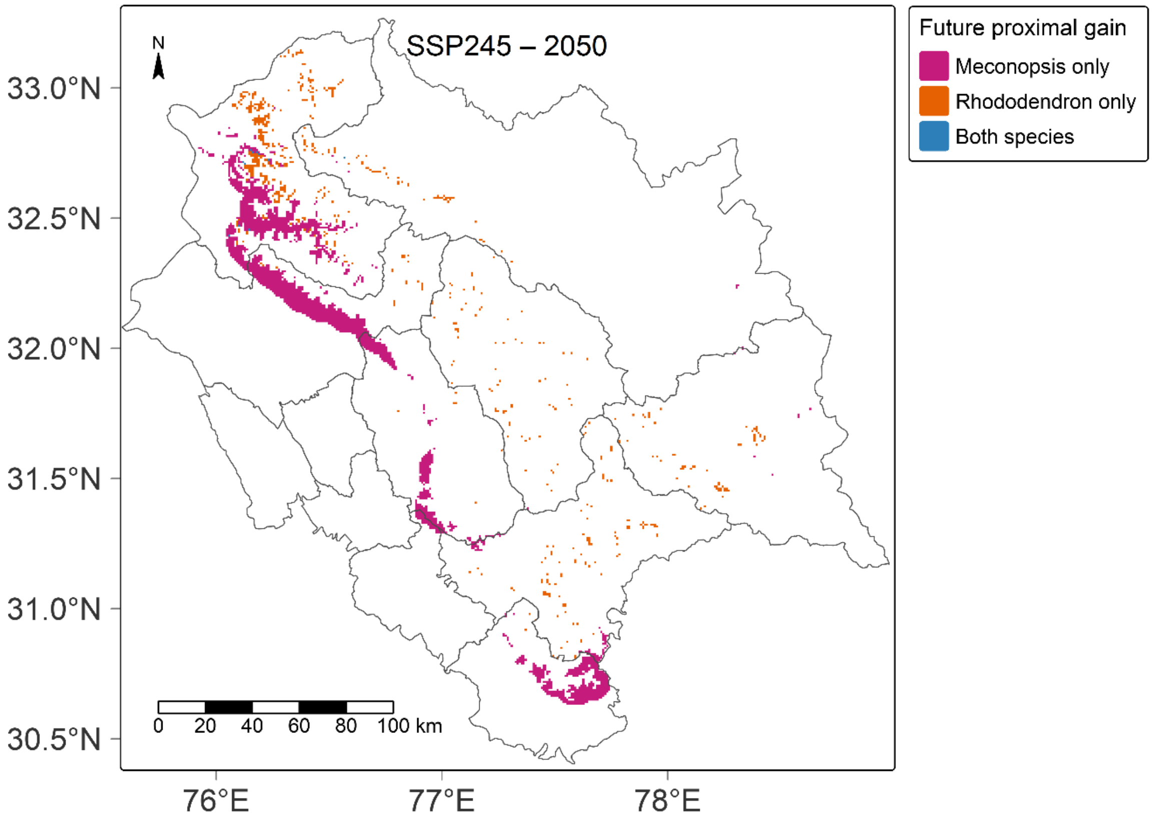
SSP245 – 2050
Future proximal gain
Meconopsis only
Rhododendron only
Both species
N
0 20 40 60 80 100 km
33.0°N
32.5°N
32.0°N
31.5°N
31.0°N
30.5°N
76°E
77°E
78°E

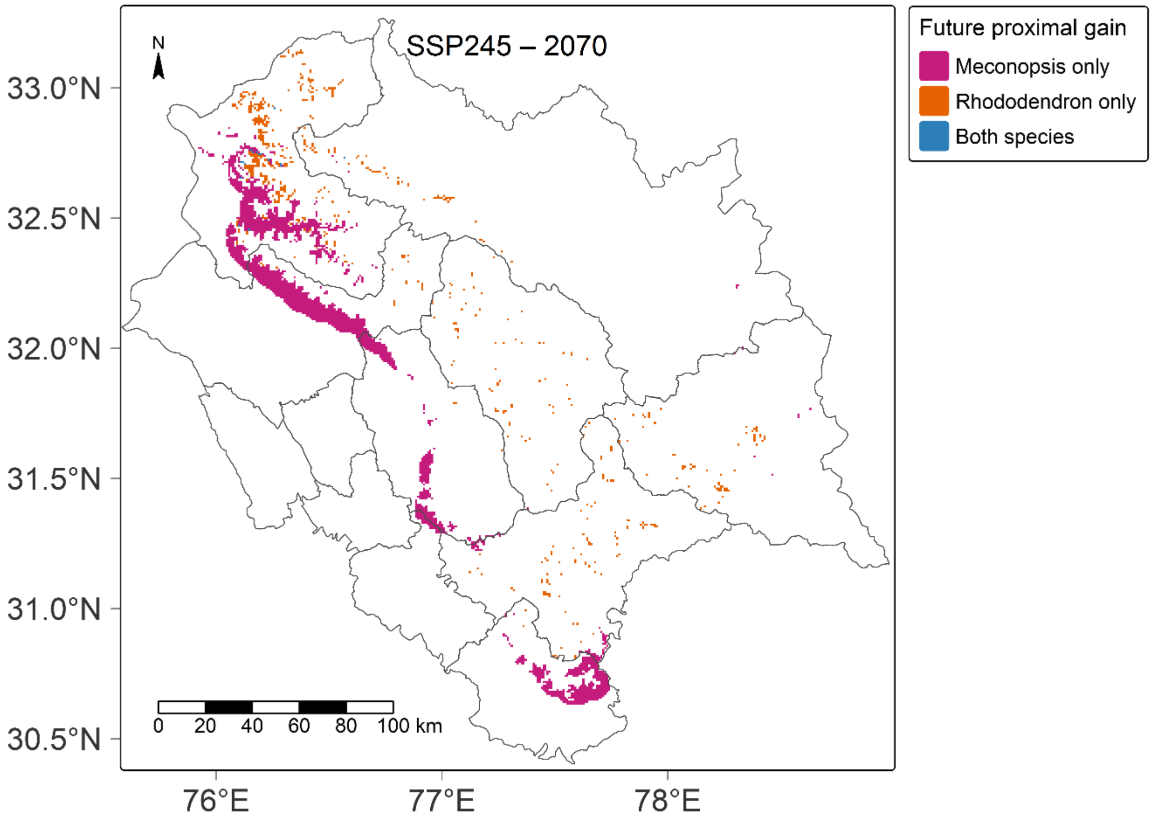
SSP245 – 2070
N
33.0°N
32.5°N
32.0°N
31.5°N
31.0°N
30.5°N
76°E
77°E
78°E
0 20 40 60 80 100 km
Future proximal gain
Meconopsis only
Rhododendron only
Both species

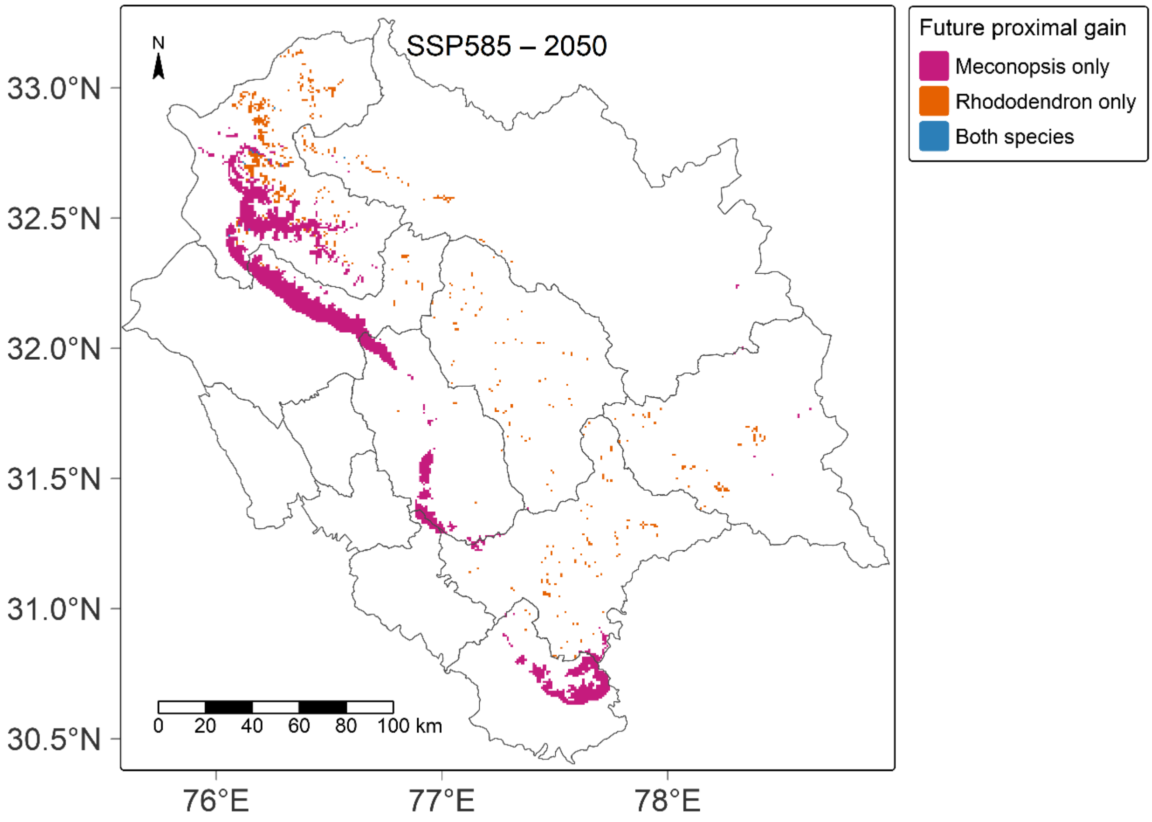


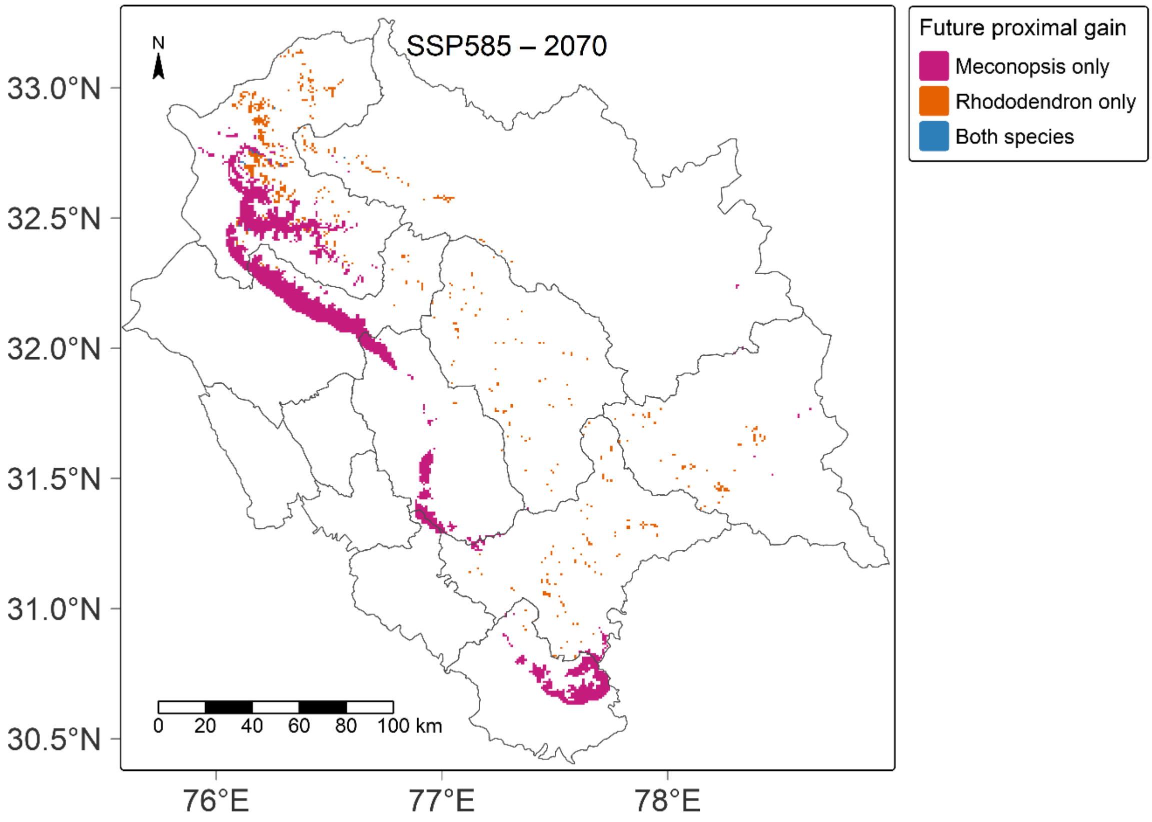


**Fig. S11**. Future proximal gain maps with suitable areas overlap for *M. aculeata and R. campanulatum*
